# Dynamic remodeling of chromatin during human mucosal-associated invariant T cell development

**DOI:** 10.64898/2026.03.17.712522

**Authors:** Marziyeh Taheri, Bowon Kim, Louis Perriman, Sedigheh Jalali, Christopher Menne, Igor E Konstantinov, Adam T Piers, Hui-Fern Koay, Stuart P Berzins, Boris Novakovic, Daniel G Pellicci

## Abstract

T cell development in the thymus is a tightly regulated process where epigenetic modifications, such as histone 3 lysine 27 acetylation (H3K27ac), play a crucial role in controlling the activation of genes. The epigenetic regulation of human mucosal-associated invariant T (MAIT) cell development is unknown; we mapped the regulatory chromatin landscape in the three developmental stages of thymic MAIT cells to identify the regulatory elements and enhancer activity involved in thymic maturation and analysed whether these chromatin dynamics are associated with the acquisition of effector programs in developing MAIT cells. Utilising cleavage under target and tagmentation (CUT&Tag), genome-wide H3K27ac profiles were generated and combined with transcriptome data from thymic MAIT cells, which revealed how developmental shifts in enhancer activity correspond to changes in gene expression. In total, 41,958 genomic regions with H3K27ac signal were identified in MAIT cells across the three development stages, of which 1,200 regions showed acetylation changes during differentiation from stage 1 to stage 3. At dynamic regions, the greatest differences were observed between stage 1 and stage 3, highlighting a progressive gain or loss of H3K27ac during MAIT cell development. Overall, MAIT cell maturation was associated with the gradual accumulation of H3K27ac at promoters and enhancers, which closely correlated with gene expression changes during development. Stage-specific enrichment of H3K27ac was observed at key transcription factor gene loci involved in MAIT cell development, including *ZBTB16* (PLZF), *EOMES*, *RUNX3*, *NFATC2*, *FOXO1*, *TGIF1*, *IRF1*, and *MAF* genes. Epigenetic remodelling was also observed at cytokine and cytokine receptors (*IL7R, IL18R1*, *IL23R, IFNG*), chemokines and chemokine receptors (*CCL4, CCL5, CCR5, CCR9, CXCR4, CXCR6*), as well as several surface molecules with known immunological function. Our work reveals a previously uncharacterised epigenetic profile of human MAIT cells that regulates and influences their development.

## Introduction

T cell development is a complex and highly regulated process, greatly influenced by epigenetic regulation of transcription factors (1, 2). Histone post-translational modifications and DNA methylation are examples of epigenetic changes during T cell differentiation (1). Specifically, acetylation of histone 3 at lysine 27 (H3K27ac) is a well-studied epigenetic mark related to active gene transcription (2, 3). This mark is found at gene promoters and distal regulatory elements, such as enhancers, and serves as a marker of active chromatin (2). By marking open or transcriptionally permissive regions, H3K27ac influences transcriptional activation and identifies regulatory regions that contribute to the developmental programming of immune cells (2, 4–6). However, although epigenetic enrichment at active regulatory regions has been reported in Mucosal-associated invariant T (MAIT) cells (7), the dynamic role of epigenetic mechanisms during human MAIT cell development remains largely unexplored.

MAIT cells are a unique subset of unconventional T cells that make up between ∼2–5% of T cells in human blood and up to ∼20–50% of T cells in certain tissues (8–10). MAIT cells express a semi-invariant T cell receptor (TCR) (TRAV1-2–TRAJ33 in humans; TRAV1–Traj33 in mice) coupled with a restricted set of TCR beta chains (Vβ2 or Vβ13 in humans and Vβ6 or Vβ8 in mice) (10, 11). This semi-invariant TCR enables MAIT cells to recognise microbial vitamin B metabolites presented by MHC class I-related protein (MR1) (8, 10, 11). Studies in MR1-deficient mice and germ-free mice revealed that the lack of MR1 molecules or riboflavin-producing commensal microorganisms leads to a significant reduction in MAIT cell frequency due to their impaired development (8, 12, 13). In mice, MAIT cell development occurs in the thymus following recognition of MR1 expressed by CD4^+^CD8^+^ double-positive (DP) cortical thymocytes (8–10, 14). Ligation of MR1 is important in both the initial commitment of T cells toward the MAIT cell lineage and in supporting further functional and phenotypic maturation of MAIT cells within the thymus (9, 11). Commensal bacteria also play a crucial role in MAIT cell expansion and full effector differentiation in the periphery (13, 15). Further, MAIT cells can also be activated by cytokines, including IL-12 and IL-18 (16). Despite increasing knowledge of the transcriptional programs regulating MAIT cell development (8, 9, 17, 18), the epigenetic mechanisms controlling these changes remain largely unknown. A key challenge in studying epigenetic regulation in MAIT cells is their relatively low frequency in the thymus (8, 9, 19), which has historically limited genome-wide chromatin analyses. Traditional epigenomic technologies such as ChIP-seq require large numbers of cells, making them difficult to apply to rare immune populations (20). Nevertheless, advances in MAIT cell identification have enabled the isolation of defined developmental subsets for further investigation. The use of MR1-5-OP-RU tetramers enabled the identification and isolation of three distinct developmental subsets of MAIT cells in both human and mouse thymus (8, 10). In mice, multiple factors are crucial for the transition between stages. For example, miRNAs are needed for the transition from stage 1 to stage 2, and the transcription factor PLZF, in combination with IL-18, for progression from stage 2 to stage 3 (8, 9, 21, 22). In humans, three stages of thymic MAIT cell development can be delineated with the markers CD27 and CD161(8). Stage 1 human MAIT cells are defined as CD27^−^CD161^−^, stage 2 MAIT cells are CD27^+^CD161^−^ and stage 3 MAIT cells are CD27 ^lo/^⁺CD161^+^ (8). While thymic stage 3 cells closely resemble circulating MAIT cells, additional extrathymic changes are needed for human MAIT cells to become fully mature (8, 9). In our previous work, we employed RNA-sequencing to compare the transcriptional changes of human thymic MAIT cells as they developed from stage 1 to stage 3, revealing 625 differentially expressed genes with notable changes to key transcription factors, and genes that regulate cell function (9). To better understand how genes are regulated during human MAIT cell development, we have now enriched thymus samples for MAIT cells and isolated the three distinct development stages for epigenetic analysis. Here, we explored how the epigenetic mark, H3K27ac, influences MAIT cell differentiation through the three stages of thymic development. The low cell input epigenomic technique cleavage under targets and tagmentation (CUT&Tag) was used, which is essential for analysing relatively rare populations of cells (23). Our work identifies a distinct H3K27ac epigenetic signature associated with human MAIT cell development, and combined with transcriptomic data, we provide a map of chromatin regions that are involved in gene regulation during this process.

## Results

In our previous studies, we defined a three-stage developmental pathway for MAIT cells, based on the differential expression of CD27 and CD161 (8, 9). These stages are characterised by distinct transcriptional profiles, which highlight dynamic changes in gene expression as MAIT cells differentiate in the thymus (9). Specifically, extensive differences were observed as cells matured in the expression of transcription factors and in molecules that regulate function, differentiation and migration, suggesting that MAIT cell development is a tightly regulated process (9). Accordingly, we hypothesised that epigenetic remodelling, particularly changes in H3K27ac, regulates gene expression in MAIT cells as they develop in the thymus. To assess this, we employed CUT&Tag epigenomic profiling to assess the role of H3K27 acetylation during MAIT cell development. Thymic MAIT cells were identified amongst human thymocytes based on staining with human MR1-5-OP-RU tetramer in combination with TRAV1-2 (Vα7.2) TCR expression. The full gating strategy of MAIT cells is shown in Supplementary Figure 1. To ensure that adequate numbers of all three thymic stages of human MAIT cells could be obtained for epigenetic analysis, thymocytes were first enriched using magnetic cell separation (MACS) technology with magnetic beads (Figure 1). MR1-5-OP-RU tetramer^+^ cells were enriched to isolate human stage 1 and 2 MAIT cells, while CD161^+^ thymocytes were separately enriched to isolate human stage 3 MAIT cells.The use of MR1-5-OP-RU tetramer leads to preferential enrichment of immature stage 1 and stage 2 MAIT cells, while the additional use of CD161 enables greater enrichment of mature stage 3 MAIT cells (Figure 1). Following MACS enrichment, cells were FACS-sorted, and MAIT cells were defined as stage 1: CD27^−^CD161^−^, stage 2: CD27⁺CD161^−^, and mature stage 3: CD27⁺/^lo^CD161⁺ cells (Figure 1).

**Figure 1:**
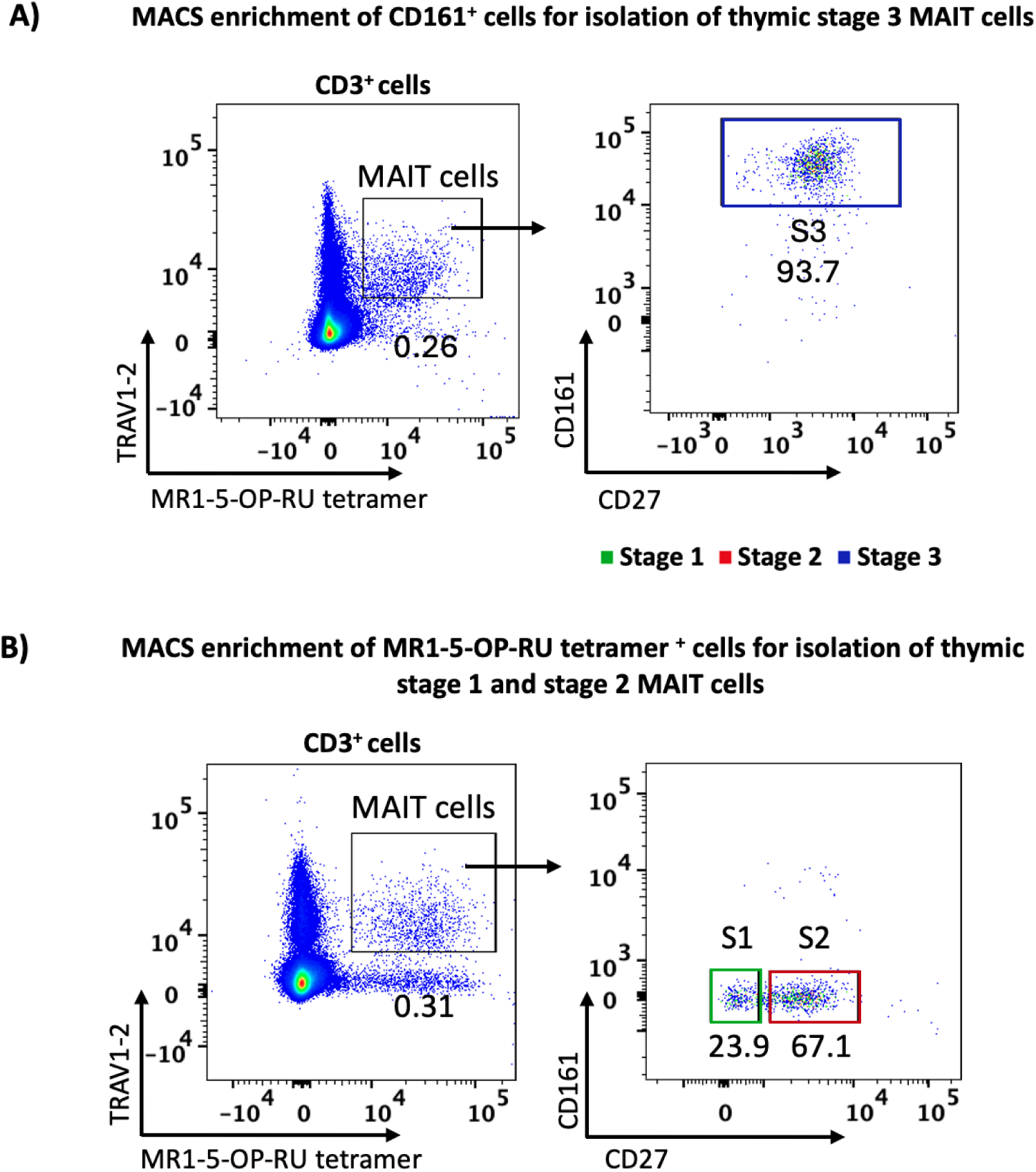
Gating strategy to define thymic MAIT cells developmental stages (stage 1, stage 2 and stage 3). **(A)** MACS enrichment of MR1-5-OP-RU tetramer^+^ cells was used to isolate stage 1 (green) and stage 2 (red) thymic MAIT cells. (**B)** MACS enrichment of CD161^+^ cells was used to isolate stage 3 thymic MAIT cells (blue). MAIT cells were identified based on MR1-5-OP-RU tetramer^+^ TRAV1-2^+^ (Vα7.2) expression. Stage 1 was defined as CD27^−^CD161^−^, stage 2 was defined as CD27^+^ CD161^−^, and stage 3 was defined as CD27⁺^/lo^CD161⁺.

### Genome-wide enrichment and distribution of H3K27ac across developmental stages of MAIT cells

Specifically, we generated genome-wide H3K27ac profiles on the three stages of MAIT cells from the human thymus. We also compared previously published bulk RNA sequencing data with the H3K27ac epigenetics data to analyse the three stages of MAIT cells (9). Assessment of H3K27ac revealed that high enrichment of acetylation occurs near transcription start sites (TSS) (Figure 2A), which is indicative of actively transcribed genes. In chromatin profiling analyses, a peak represents a genomic region where sequencing reads accumulate, showing local enrichment of the histone modification at that site. Across all three developmental stages of MAIT cells, we identified a total of 41,958 H3K27ac marked peaks (Figure 2B). 65% of these peaks were mapped to distal regulatory regions, while 35% were mapped to promoters (i.e. within 5kb of TSS) (Figure 2B). Moreover, a high percentage of peaks (86%) were detected in the open chromatin regions (Figure 2B), indicating dynamic remodelling of chromatin during MAIT cell development. From a total of 41,958 H3K27 ac peaks, 11,132 peaks were detected within 1 Mb of annotated genes (Figure 2C). Based on genomic proximity, each peak was mapped to its nearest gene, and this approach enabled us to identify 4,158 genes with at least one peak, 2,339 genes with 2 peaks, 3,758 genes with 3-9 peaks and 877 genes associated with ten or more peaks (Figure 2C).

**Figure 2.**
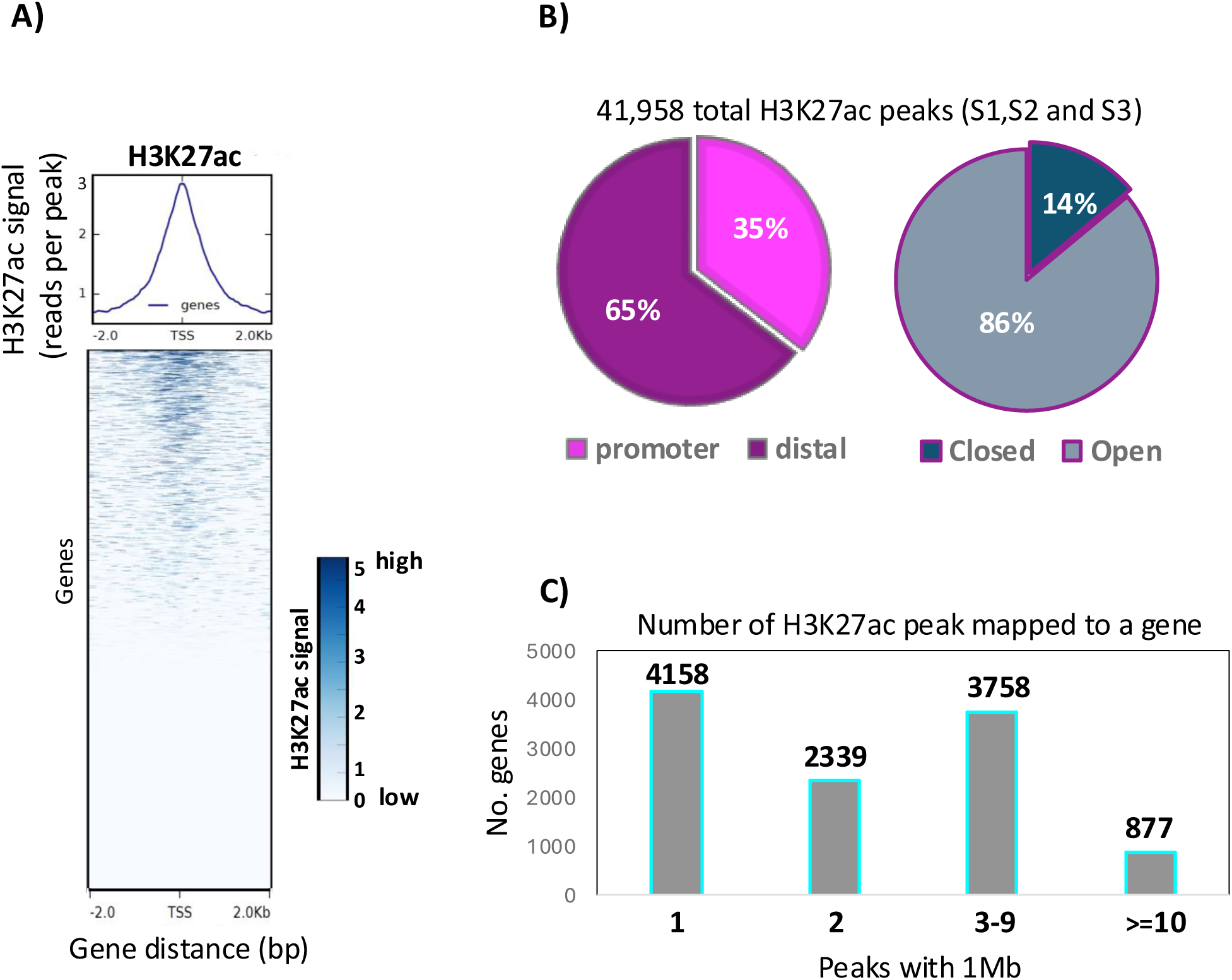
Genomic distribution of H3K27ac peaks during MAIT cell development. **(A)** Heatmap of H3K27ac signal across ±2 kb anking transcription start sites (TSSs). **(B)** Pie charts show the genomic distribution of H3K27ac peaks located in distal regulatory regions (purple), or promoter regions (pink), and in closed chromatin regions (teal) or open chromatin regions (grey). **(C)** The bar charts depict the distribution of genes based on the total number of H3K27ac peaks within ±1 Mb of TSSs.

### MAIT cell differentiation is associated with progressive remodelling of H3K27ac

To identify epigenetic changes and regulatory elements associated with MAIT cell development, we visualised the H3K27ac levels across all three developmental stages at peaks that gained or lost acetylation. Out of the total of 41,985 H3K27ac peaks, 1,263 peaks (3%) exhibited significant differential enrichment from stage 1 to stage 3 thymic MAIT cells, 1,020 peaks gained acetylation (Figure 3A), while 243 peaks lost acetylation (Figure 3B). Principal Component Analysis (PCA) based on the 2,414 dynamic peaks (5.7% of all detected peaks) revealed a clear separation of three MAIT cell developmental stages (Figure 3C). Stage 1 was most separated from stage 2 and 3 on PC1, indicating that it was most different in its H3K27ac profile, followed by the separation of stage 2 and 3 on PC2. These data support progressive chromatin remodelling during MAIT cell differentiation (Figure 3C). We also detected that the greatest differences in H3K27ac occurred when comparing stage 1 with stage 3, whereas fewer changes were observed when comparing stage 1 versus stage 2 or stage 2 versus stage 3 MAIT cells (Figure 3A and 3B). This analysis identified several hundred dynamic peaks between stage 1 and stage 3 of MAIT cell development (Figure 3D). Several genes associated with MAIT cell maturation, including *ZBTB16* (PLZF), *MAF, STAT4, NFATC2, FOXO1, EOMES, CST7* and *IL7R* (CD127), displayed increased H3K27ac in stage 3 MAIT cells (Figure 3D), which matched increased expression of these same genes in stage 3 cells, compared to stage 1 MAIT cells (9). Interestingly, H3K27ac of *RUNX1, BCL11B,* and *CCR9* was increased in stage 3 MAIT cells (Figure 3D), but had reduced gene expression in stage 3 cells (Supplementary file 1) (9). Genes such as *LEF1, SATB1, TOX2,* and *SOX4* exhibited higher H3K27ac in stage 1 MAIT cells (Figure 3D), which is consistent with transcriptomic data and supported their role in early T cell development (a full list of genes is provided in Supplementary file 1) (9).

**Figure 3:**
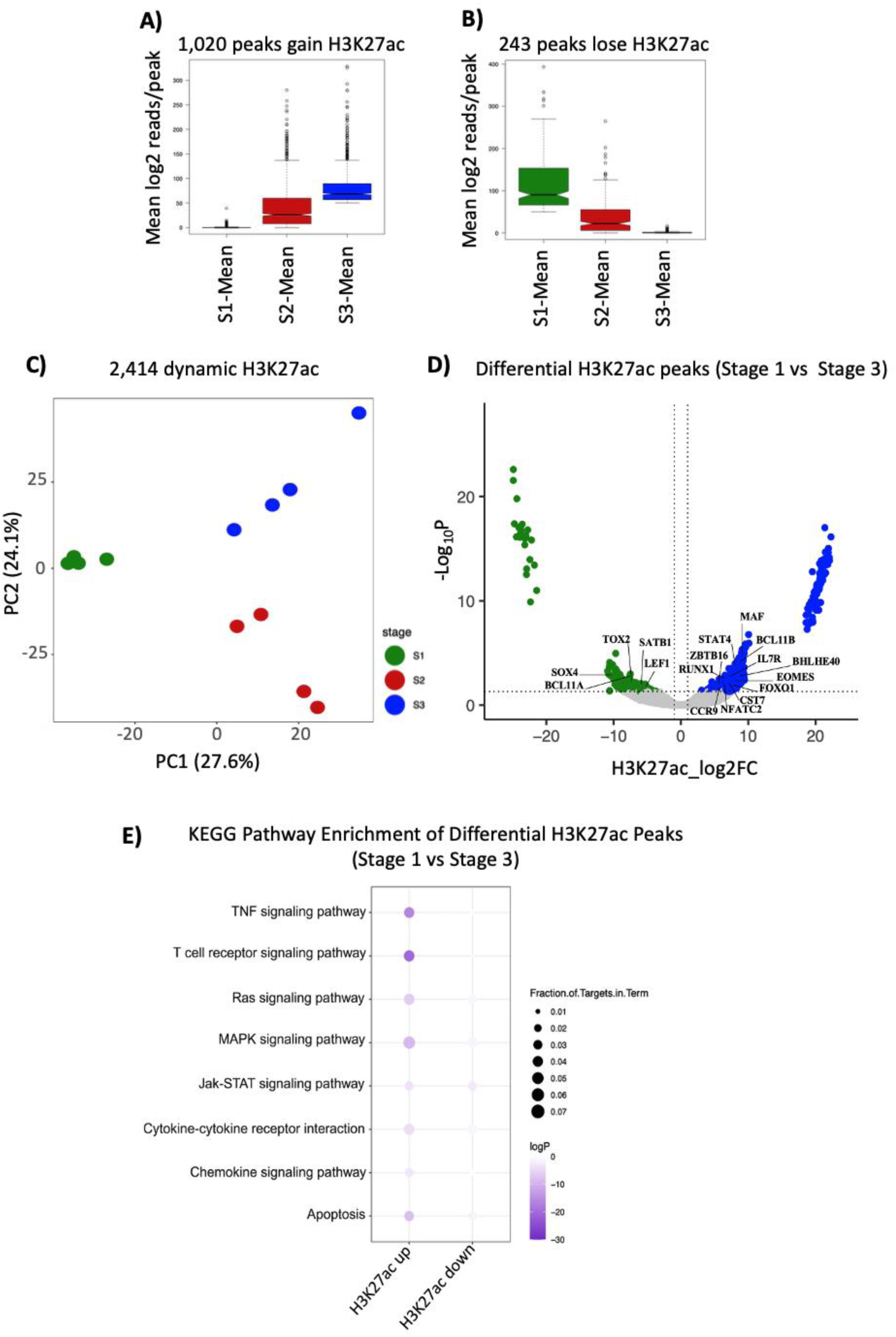
Dynamic changes in H3K27ac peaks enrichment across MAIT cell developmental stages. **(A)** Box plot displaying peaks that gained acetylation and **(B)** lost acetylation during MAIT cell development. The mean H3K27ac enrichment was calculated from four samples per stage. Each colour represents stage 1 (green), stage 2 (red) or stage 3 MAIT cells (blue). The peaks were identified using DESeq2 for significance thresholds (padj<0.05), and the boxplots visualise normalised signals (log2 reads per peak) across the three stages. **(C)** PCA plot of the differential H3K27ac profile. The first principal component (Dim 1) explains 27.5% of the variance, and the second principal component (Dim 2) explains 24.1% of the variance. Each stage included four biological replicates. **(D)** Volcano plot displaying differential H3K27ac peaks between stage 1 and stage 3 MAIT cells based on H3K27ac log₂ FC between two stages (x-axis), and −log₁₀ adjusted p-value (–log₁₀ P) from DESeq2 analysis (y-axis). Each dot represents an individual H3K27ac peak, and one representative acetylation peak per gene is shown. Blue dots show peaks with significantly higher acetylation in stage 3, and green dots show peaks with higher acetylation in stage 1 (padj < 0.05, log₂ FC > 2). Grey dots show non-significant peaks. **(E)** The bubble plot shows the enriched KEGG pathways with H3K27ac, based on significance (logP) and the fraction of genes associated with each pathway.

Pathway analysis of genes associated with differential H3K27ac between stage 1 and stage 3 thymic MAIT cells revealed significant enrichment of multiple immune pathways, indicating robust epigenetic activation of these pathways (Figure 3E). Pathways that gained H3K27ac, including cytokine–cytokine receptor interaction and JAK–STAT signalling, were accompanied by increased acetylation at loci encoding *IL18R1*, *IL18RAP*, *IL23R*, *IL2RB*, *IL7R* and *STAT4* (Figure 3E). T cell receptor, Ras, and MAPK signalling pathways were associated with increased acetylation at loci encoding transcriptional regulators, including *ZBTB16* (PLZF), *EOMES*, *RUNX3* and *FOXO1* (Figure 3E). Enrichment of the TNF signalling pathway was associated with H3K27ac changes at downstream signalling molecules such as *NFKB1*, while chemokine signalling pathway enrichment was associated with migration-related genes (*CCR5*, *CXCR6*, *CCL4*, *CCL5*) and *S1PR1*. Enrichment of the apoptosis pathway was supported by epigenetic changes at genes involved in lymphocyte survival and cytotoxicity, including (*BCL2*, *GIMAP* family members, *GZMA* and *PRF1*) (Figure 3E). Collectively, these changes suggest that enhancer activation at these regions may contribute to the functional priming of MAIT cells during thymic maturation.

### H3K27ac remodelling and analyses of MAIT cell related genes during maturation from stage 1 to stage 3

To explore epigenetic changes of key genes during MAIT cell development, we first counted the dynamic H3K27ac peaks detected across the three MAIT cell stages. A total of 3,081 genes had a H3K27ac peak within 1Mb of their TSS, ranging between 1 to 15 peaks per gene (Figure 4A and Supplementary file 3). Transcription factors (TFs) thought to be critical for MAIT cell development were enriched in regions that contained at least 3 to 15 H3K27ac peaks, reflecting the dynamic control of their expression through multiple promoters and distal elements (Figure 4A). Remarkably, *BCL11B* and *ZBTB16* (PLZF) had 15 and 10 peaks, respectively, and ranked among the most acetylated and epigenetically modified transcription factors during MAIT cell development. Other genes for transcription factors, including *BHLHE40*, *FOXO1*, *ETS1, ID2*, *MAF, EOMES, RUNX3, STAT4* and *TCF7,* also contained three or more peaks, suggesting an important role for these genes in regulating MAIT cell development (Figure 4A and supplementary file 3). To understand how H3K27ac compares to gene expression in developing MAIT cells from human thymus, we revisited our published RNA-sequencing data that described differentially expressed genes (DEGs) in thymic MAIT cells that matured from stage 1 to stage 3 (9). *MAF, BHLHE40, EOMES, STAT4, RUNX3, ZBTB16* (PLZF)*, FOXO1* and *ID2* exhibited increased gene expression as cells matured to stage 3 (Figure 4B). Whereas, *ETS1, TCF7, FLI1* and *LEF1* showed reduced RNA expression as cells matured to stage 3 (Figure 4B). Although *BCL11B* exhibited the highest number of H3K27ac marks, indicating substantial epigenetic remodelling, its RNA expression decreased during maturation (Figure 4 A, B), demonstrating that an increase in chromatin acetylation is not always accompanied by enhanced gene expression. To better understand how chromatin changes may influence transcription factor binding during MAIT cell development, we examined the presence of transcription factor binding motifs within dynamic H3K27ac regions (Figure 4C). As displayed schematically, transcription factor motifs are distributed at distal regulatory elements with changing histone acetylation (Figure 4C). Motif enrichment analysis revealed that binding motifs for *ZBTB16* (PLZF), *FOXO1, FLI1, EOMES, BCL11B*, *ERG, RUNX3*, *STAT4, LEF1, SOX4*, *MAF, ID2, TCF7* and *BHLHE40* were dominant in regions that gained acetylation, whereas a lower frequency of binding motifs was detected in the regions that lost acetylation (Figure 4C). We used several published PLZF binding motifs (24–26) and scanned for those regions. Our analysis revealed a complex PLZF binding signature associated with increased H3K27ac at regulatory regions bound by PLZF. Increased H3K27ac was also observed at sites where PLZF co-localised with the histone methyltransferase *EZH2*, as reported by Koubi et al (24) (Figure 4C). Specifically, peaks containing two motifs for *PLZF-EZH2* and *BCL11B* showed higher H3H27ac signal in comparison with peaks containing one or no motifs (Figure 4D). Many of these transcription factors, including *ZBTB16*, *EOMES*, *MAF*, *RUNX3* and *ID2*, are known regulators of unconventional T cell differentiation, suggesting that progressive enhancer activation at their loci may reinforce lineage-specific transcriptional networks during MAIT cell maturation.

**Figure 4:**
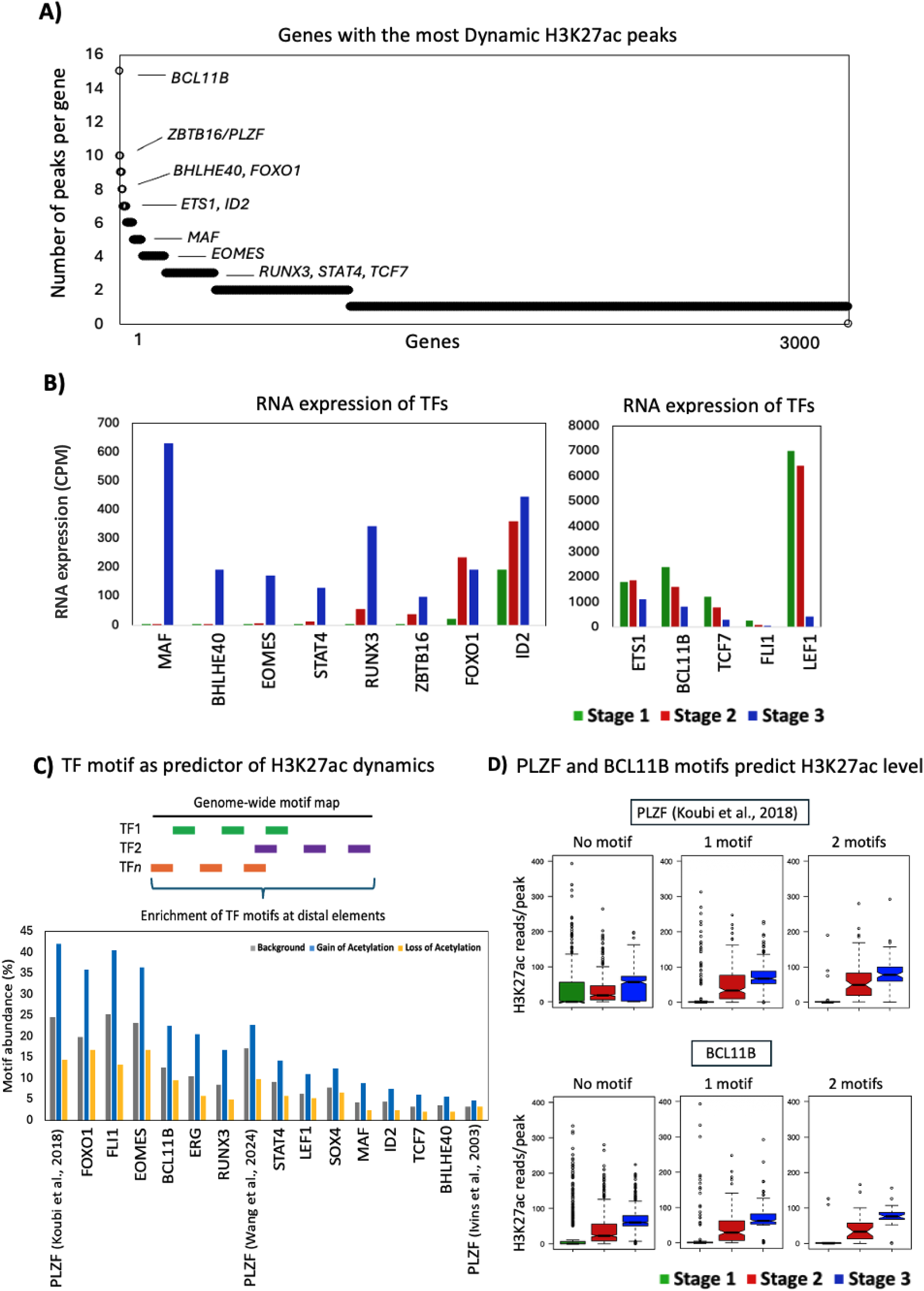
H3K27ac profiling and motif enrichment analyses in MAIT cell development. **(A)** The dot plot of the top genes with the highest number of differential H3K27ac peaks across the three stages of human thymic MAIT cells. Peaks were identified using DESeq2 for significance thresholds (padj<0.05). Peaks were annotated to nearby genes using a genomic annotation tool **(B)** The bar charts show the normalised RNA expression levels (count per million, CPM) of genes involved in MAIT cell development. **(C)** Bar chart showing enrichment of gene motifs at dynamic H3K27ac in enhancer regions with gain of acetylation (blue), loss of acetylation(yellow), and background regions (grey). **(D)** Boxplots showing H3K27ac signal (reads/peak) for region with no motif, one or two motifs for PLZF-EZH2 (top) and BCL11B (bottom). Each colour represents stage 1 (green), stage 2 (red) or stage 3 MAIT cells (blue). Statistical comparisons between motif groups (0–2 motifs) were performed within each developmental stage using t-tests. P values < 0.05 were considered statistically significant.

### Expression level of transcription factors involved in MAIT cell differentiation

Next, we examined for overlap between gene RNA expression and H3K27ac enrichment across individual regulatory peaks for transcription factors identified as important for MAIT cell development, including *ZBTB16* (PLZF), *BCL11B*, *FOXO1* and *LEF1* (Figure 5). *ZBTB16* displayed an increase in gene expression together with enhanced H3K27ac across several peaks as cells matured from stage 1 to stage 3 (Figure 5A). While *BCL11B* gene expression decreased as cells developed from stage 1 to stage 3, H3K27ac peaks associated with *BCL11B* were highest in stage 3 cells compared to stage 1 cells (Figure 5B). RNA expression levels of *FOXO1* were increased in stage 3 compared to stage 1 MAIT cells, while its H3K27ac levels showed a mixed pattern, with 8 out of 10 peaks gaining H3K27ac, while 2 peaks showed decreased acetylation in stage 3 cells compared to stage 1 MAIT cells (Figure 5C). Conversely, *LEF1* showed reduced gene expression as cells matured from stage 1 to stage 3 MAIT cells, but its H3K27ac profile was variable across the 5 peaks, with some losing acetylation as cells matured to stage 3, while others gained acetylation (Figure 5D). Accordingly, key transcription factors involved in MAIT cell development display distinct H3K27ac patterns across their regulatory regions, indicating that chromatin remodelling contributes to their regulation during MAIT cell development.

**Figure 5:**
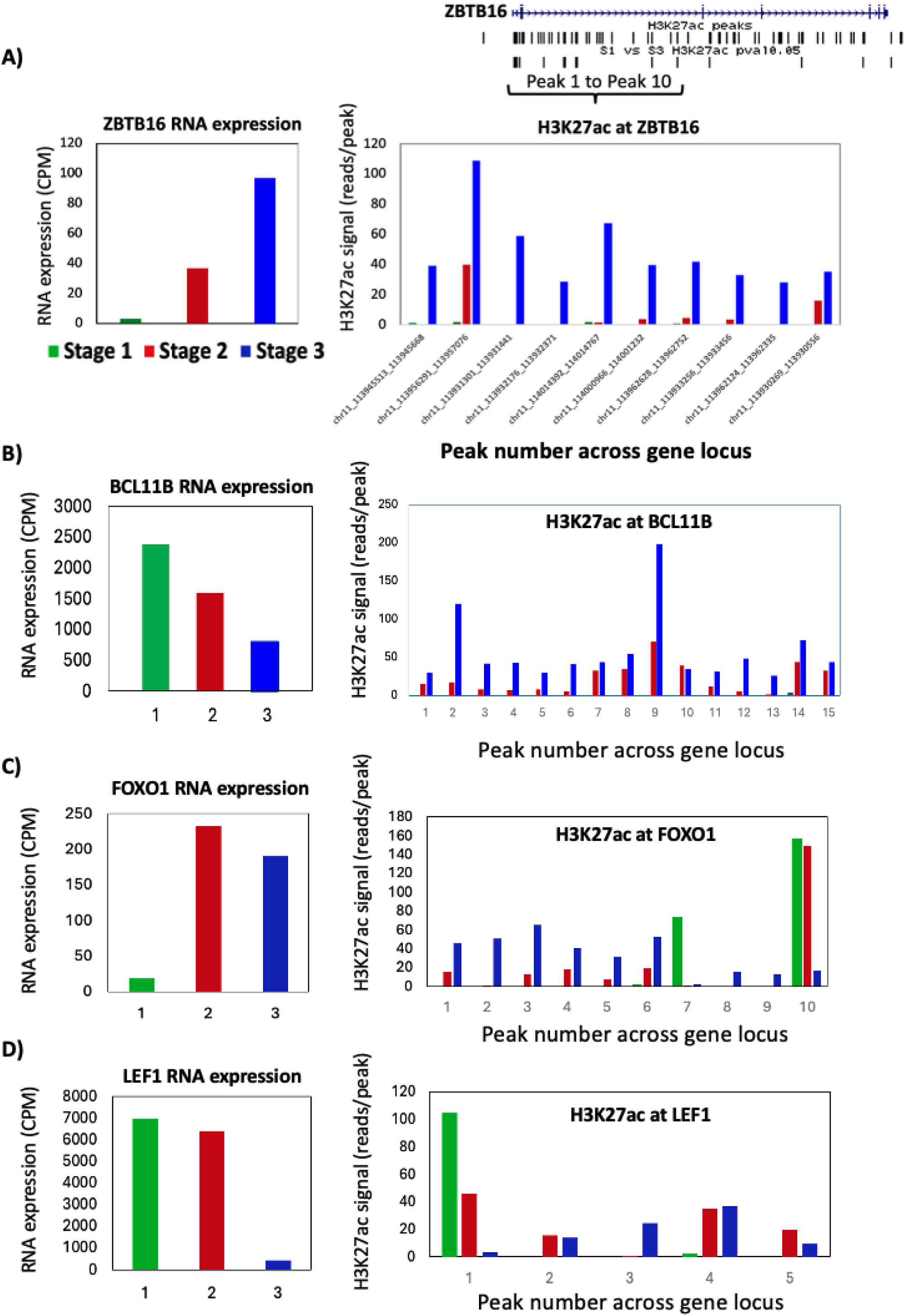
Correlation between H3K27ac enrichment and gene expression of ZBTB16, BCL11B, FOXO1 and LEF1 during MAIT cell development. **(A-D)** Bar charts show the side-by-side correlation between RNA expression levels (left) and H3K27ac signals (right) across developmental stages of human thymic MAIT cells for (A) *ZBTB16*, (B) *BCL11B*, (C) *FOXO1* and (D) *LEF1.* Each colour represents one stage: stage 1 (green), stage 2 (red), and stage 3 (blue). RNA expression is based on the mean of counts per million (CPM), while individual acetylation peak locations across each gene locus are based on H3K27ac signal intensity (reads per peak). Statistical analysis was performed using DESeq2 for differential peak enrichment (*padj* < 0.05), and CPM normalisation was applied to compare gene-expression levels across stages.

### Dynamic H3K27ac strongly correlates with gene expression by developing MAIT cells

We compared stage 1 with stage 3 based on the correlation between the number of H3K27ac peaks and transcript levels (Figure 6A). This analysis showed that 538 peaks gained acetylation on stage 3 MAIT cells and were associated with nearby genes that also exhibited higher transcript levels (Figure 6A). Conversely, 123 peaks lost acetylation on stage 3 MAIT cells that were accompanied by reduced RNA expression (Figure 6A). Only a small number of peaks exhibited a mixed pattern. For example, 188 gained acetylation, which was accompanied by reduced gene expression, while 37 lost acetylation but had increased gene expression (Figure 6A). Using our published MAIT cell transcriptome dataset (9), we further examined this relationship across promoter (within 2.5 kb of the TSS) and distal regulatory regions (>5 kb from the TSS) throughout the three developmental stages (Figure 6 and supplementary tables 1-10). Overall, H3K27ac levels were generally positively associated with gene expression changes during thymic MAIT cell maturation from stage 1 to stage 3.

**Figure 6:**
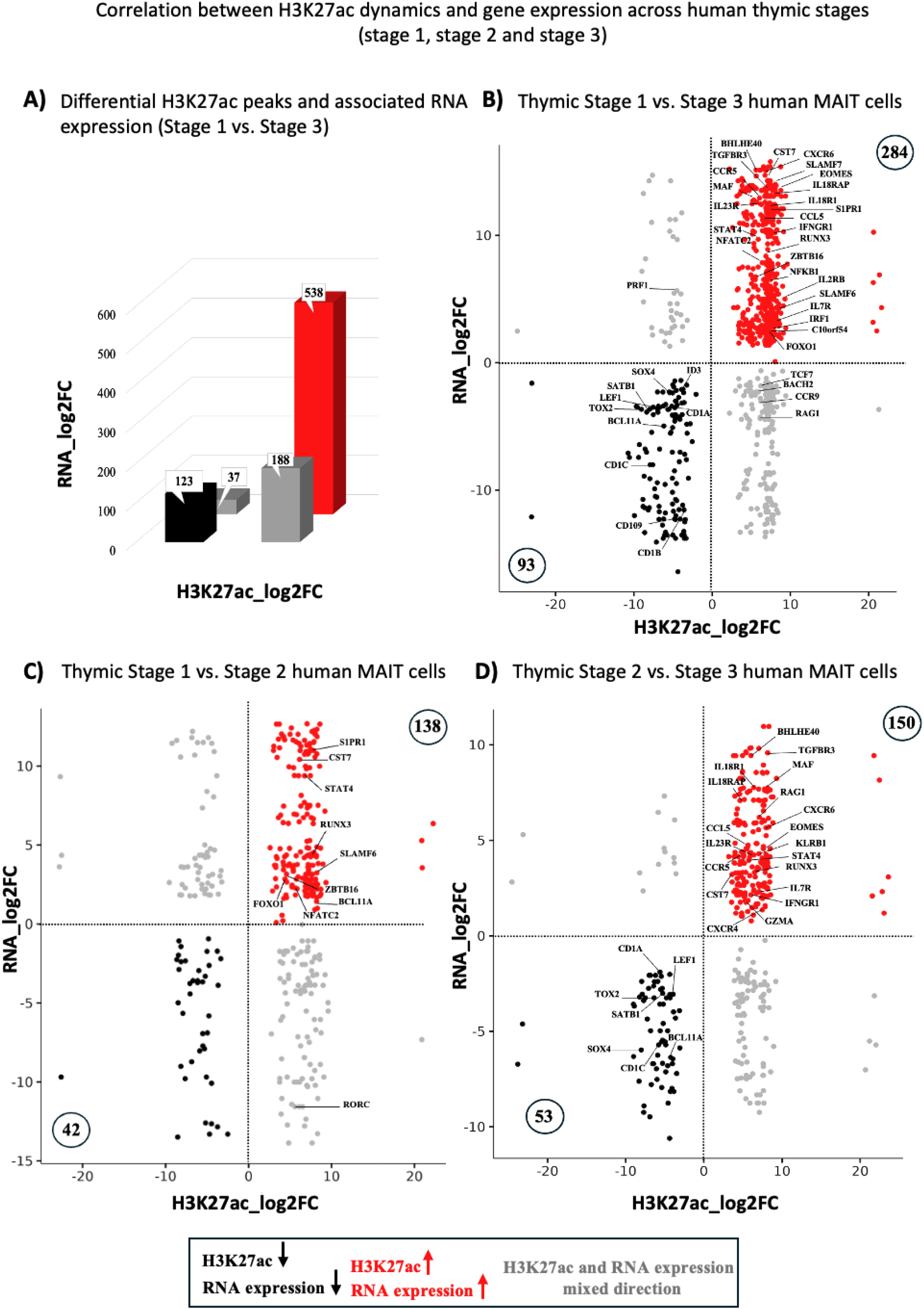
Correlation between H3K27ac enrichment and gene expression during MAIT cell development. **(A)** Bar chart showing the correlation between H3K27ac peaks and transcript levels comparing stage 1 to stage 3 MAIT cells. Each bar represents the number of acetylation peaks associated with nearby genes in correlation with RNA expression changes. Bar charts (red) show increased both acetylation and RNA expression, (black) show decreased acetylation and RNA expression, and (grey) show a mixed trend. Scatter plot showing the correlation between H3K27ac (log2 fold change, x-axis) and gene expression (CPM log2FC, y-axis) across thymic developmental stages of MAIT cells (**B)** comparing stage 1with stage 3, **(C)** stage 1 with stage 2 and **(D)** stage 2 with stage 3. Each dot represents a peak, and only one representative acetylation peak per gene is shown. Red dots show genes with increased RNA expression and H3K27ac levels, black dots show genes with decreased RNA expression and H3K27ac levels, and grey dots indicate genes that show increased RNA expression but not H3K27ac, or increased H3K27ac but not RNA expression. Pearson correlation analysis was used to assess the association between H3K27ac and RNA expression. Each stage included four biological replicates. H3K27ac peaks were identified using DESeq2, and gene expression analyses were performed using R. *P* < 0.05 is considered significant.

A total of 284 genes displayed a significant increase in both acetylation and gene expression (common up) on stage 3 cells compared to stage 1 MAIT cells (Figure 6B and supplementary table 2). Whereas 93 genes were decreased in both acetylation and gene expression on stage 3 cells compared to stage 1 MAIT cells (Figure 6B and supplementary table 1). Comparing stage 1 with stage 2 MAIT cells, 138 genes were significantly upregulated for both acetylation and gene expression by stage 2 cells (Figure 6C and Supplementary Table 5). Whereas, 42 genes were significantly downregulated in expression, marked by reduced H3K27ac by stage 2 cells compared to stage 1 MAIT cells (Figure 6C and Supplementary Table 4). When comparing stage 2 with stage 3 cells, 150 genes showed an increase in both acetylation and gene expression in stage 3 cells (Figure 6D, Supplementary Table 8), and 53 genes were commonly downregulated for acetylation and gene expression (Figure 6D, Supplementary Table 7). Accordingly, the greatest changes in both acetylation and transcript levels occurred between stage 1 and stage 3 thymic MAIT cells, supporting the predicted developmental trajectory of these cells (Figure 6B) (9, 11).

To better understand the genes that regulate thymic development of human MAIT cells, we examined genes showing a significant and clear trend across developmental stages, including those that were consistently upregulated, downregulated or showed mixed patterns between transcript levels and H3K27ac (the full list of genes is provided in supplementary table 1-10). The transcription factors *ZBTB16*, *EOMES, MAF, STAT4, NFATC2, NFKB1, FOXO1, IRF1, RUNX3,* and *BHLHE40* displayed significant increases in both transcript levels and H3K27ac enrichment from stage 1 to stage 3. (Figure 6B and Supplementary Table 2). In addition, members of the Ikaros family of transcription factors showed dynamic regulation as thymic MAIT cells developed (Supplementary tables 2 and 10, Supplementary file 1). Specifically, *IKZF1* (Ikaros) and *IKZF2* (Helios) showed increased H3K27ac despite the absence of detectable transcript levels. Specifically, *IKZF1* showed increased H3K27ac from stage 1 to stage 2, while *IKZF2* showed increased H3K27ac as cells mature from stage 1–2 and from stage 1–3. In contrast, *IKZF3* (Aiolos) showed both increased H3K27ac and gene expression from stage 1 to stage 3 (Supplementary file 1). Krüppel-like factors (KLF) are a family of zinc-finger transcription factors that have diverse roles in cell differentiation, activation, and development (27, 28) and are also highly regulated during MAIT cell development. Genes encoding *KLF3* and *KLF6* showed increased transcription together with enrichment for H3K27ac, when comparing stage 1 to stage 3 cells and during transition from stage 2 to stage 3 (Supplementary tables 2 and 8). Several members of the GTPase of the immunity-associated protein (GIMAP) family, which play important roles in regulating cell survival, appear important for MAIT cell development. *GIMAP5*, *GIMAP6* and *GIMAP7* were marked by increased H3K27ac, resulting in higher gene expression as MAIT cells matured to stage 3 (Supplementary tables 2, 5 and 8). *GIMAP4* and *GIMAP8* gene expression also changed as MAIT cells developed, although H3K27ac of peaks associated with these genes were not significantly different between the three stages (Supplementary file 1). The anti-apoptotic protein BCL-2, which also regulates cell survival (29), displayed increased H3K27ac at distal regulatory elements, whereas *BCL2* gene expression decreased as MAIT cells matured (Supplementary tables 3 and 9).

Genes that encode cytokines, cytokine receptors or immune regulatory surface molecules, including *TGFBR3, CST7, IL18RAP, IL18R1* (CD218)*, IL23R, SLAMF6* (CD352)*, SLAMF7* (CD319)*, IFNGR1, IFNG* (IFN-γ), *IL2RB* (CD122) and *IL7R* (CD127) were upregulated as cells matured to stage 3 and this corresponded with increased H3K27ac peaks associated with these genes (Figure 6B and supplementary table 2). These data reveal that as MAIT cells mature, they acquire greater effector functions during development and are likely to become more responsive to cytokine-driven activation. The transcription factors *TOX2, LEF1, SATB1, SOX4, ID3* and *BCL11A* and surface molecules *CD1A, CD1B* and *CD1C*, and *CD109* showed reduced H3K27ac enrichment and RNA expression as cells matured from stage 1 to stage 3. (Figure 6B and supplementary table 1). Notably, our analysis also identified several genes that showed distinct differences between transcript level and H3K27ac. For example, the *PRF1* (perforin) gene displayed increased gene expression in stage 3 MAIT cells, but had reduced H3K27ac levels compared to stage 1 cells (Figure 6B). Additionally, genes such as *TCF7, BACH2, CCR9* and *RAG1* displayed increased H3K27ac as cells matured to stage 3, while transcript levels of these genes decreased during development (Figure 6B and supplementary table 3). *RORC* (RORγt) also showed a mixed regulatory pattern; H3K27ac increased when comparing stage 1 to stage 2 MAIT cells. However, gene expression significantly decreased as MAIT cells transitioned from stage 1 to stage 2 (Figure 6C and supplementary table 6). Analysis of genes encoding chemokines and chemokine receptors associated with cell migration, including *CCL4, CCL5, CCR5* and *CXCR6*, showed increased H3K27ac and higher gene expression in stage 3 compared to stage 1 MAIT cells (Supplementary Table 2). These genes were also upregulated as MAIT cells transitioned from stage 2 to stage 3 (Supplementary Table 8). *S1PR1* is a key regulator of thymic egress (30, 31), which also exhibited increased H3K27ac and gene expression as cells matured from stage 1 to stage 3 (Figure 6B, 6C and Supplementary tables 2 and 5). Some cytotoxic-related genes, including *GZMA* (Granzyme A), as well as the surface marker (CD161) and the migration related chemokine receptor *CXCR4*, were upregulated at the transcript levels and showed increased acetylation only during the transition from stage 2 to stage 3 (Figure 6D and supplementary table 8) (32). Taken together, our analyses identified regulatory genes with dynamic epigenetic changes that may play critical roles in MAIT cell differentiation, function and migration, including a large number of transcription factors involved in T cell development (33).

We analysed a total of 1,500 validated transcription factors, of which 188 showed dynamic changes in mRNA expression between stages (Supplementary file 2). Among these, 148 transcription factors were associated with H3K27ac peaks, containing a total of 947 peaks. Of these 148 transcription factors, 75 were expressed at the RNA level and showed dynamic H3K27ac peaks (229 peaks in total), suggesting that gene expression of these transcription factors is accompanied by acetylation changes during MAIT cell maturation (Supplementary table 10 and Supplementary file 2). Notably, some transcription factor associated peaks displayed sharp changes in acetylation, either increasing or decreasing during MAIT cell development, while others changed more gradually (supplementary table 10). Transcription factors with sharp dynamic acetylation changes are more likely to play important regulatory roles during MAIT cell development. For example, *ZBTB16*, *EOMES, RUNX3, NFATC2, FOXO1, TGIF1, IRF1,* and *MAF* sharply increased during maturation to stage 3, while *SMAD1, MEF2D* and *RORB* were sharply decreased as cells matured to stage 3 (supplementary table 10). Slower acetylation dynamics were observed at several transcription factor loci, including (*BHLHE40*, *KLF6*, *FOSB*, *RORA* and *SMAD7*), which gradually increased, and (*TOX2*, *SATB1* and *SOX4*), which gradually decreased, as cells matured from stage 1 to stage 3 (Supplementary Table 10).

### Dynamic H3K27ac at promoters and distal elements correlates with dynamic gene expression across MAIT cell stages

We also separately analysed promoter and distal associated genes with epigenetic changes (Supplementary Figure 2). This analysis was performed to determine whether H3K27ac enrichment at promoter and distal regulatory elements, including enhancers, showed distinct relationships with gene expression during MAIT cell development. A positive correlation between changes in H3K27ac signals and gene expression was observed in both promoter and distal regions when comparing stage 1 with stage 3 MAIT cells (Supplementary Figure 2A). Comparing stage 2 and stage 3 MAIT cells showed an intermediate positive correlation, particularly at distal regulatory regions (Supplementary Figure 2B), while comparing stage 1 with stage 2 cells revealed a positive correlation only for distal peaks (Supplementary Figure 2C). Together, these findings highlight that both promoter and distal regulatory regions contribute to transcriptional changes during MAIT cell development, and stronger correlations are observed at later stages of maturation. This pattern likely indicates progressive enhancer activation during thymic maturation associated with the acquisition of effector programs before MAIT cells exit the thymus.

## Discussion

Epigenetic regulation controls T cell lineage commitment, differentiation and acquisition of effector functions (1). While the transcriptional landscape of MAIT cell development has been examined (9, 11, 17, 18), little is known about the epigenetic modifications that underpin these transcriptional changes in MAIT cells and other unconventional T cells in the human thymus. Epigenetic profiling may also help resolve instances where gene expression and protein levels do not align, and may distinguish genes involved in stable lineage commitment from those that are transiently expressed during development. In addition, epigenetic marks at regulatory elements may help identify genes that are functionally important for MAIT cell differentiation beyond analysis of transcriptional abundance. In this study, we show that MAIT cell development is accompanied by extensive remodelling of active chromatin landscapes, providing an epigenetic framework that helps explain transcriptional and functional maturation of MAIT cells in the thymus. Our data also reveals that while H3K27ac enrichment marks active chromatin regions, it does not always indicate direct regulation of gene expression, but rather highlights candidate regulatory elements associated with MAIT cell maturation.

Developing MAIT cells showed significant enrichment of H3K27ac across 3,081 genes, involved in transcriptional regulation, cytokine signalling, immune regulation, lymphocyte survival, cytotoxic function and chemokine receptor pathways. Notably, many transcription factors known to regulate MAIT cell lineage commitment and maturation in the thymus showed strong epigenetic enrichment (33, 34). For example, loci encoding *ZBTB16* (PLZF), *BCL11B, EOMES*, *IKZF1*, *IKZF2*, *MAF*, *STAT4*, *RUNX3*, *BACH2*, and *BHLHE40* displayed extensive H3K27ac enrichment during MAIT cell development, suggesting that the functional programming of MAIT cells is coordinated through chromatin-level regulation of key transcriptional factors. Consistent with this, previous studies have shown that *BCL11B* binds H3K27ac-enriched regulatory regions linked to TCR signalling genes in human MAIT cells, supporting enhancer-level control of MAIT transcriptional programs (7). In line with this, we observed extensive H3K27ac enrichment at the BCL11B locus but reduced transcript levels during maturation, suggesting that *BCL11B* gene expression may be regulated by additional developmental signals or reflect a primed chromatin state that is maintained despite reduced transcription. Notably, many of these transcription factors are also differentially expressed in γδ T cells and NKT cells within the thymus in both humans and mice (9–11, 35–37), suggesting that the epigenetic regulation observed for MAIT cells is likely shared with the development pathways of other subsets of unconventional T cells.

PLZF is a key lineage transcription factor regulating the innate-like characteristics of unconventional T cells (9, 10, 34, 38–40). Ikaros functions upstream of PLZF, where PKD-dependent activation of Ikaros promotes PLZF transcription, and disruption of this pathway leads to defective NKT cell development in mice. Ikaros (*IKZF1*) and Helios (*IKZF2*) are key members of the Ikaros zinc-finger transcription factor family, with Helios being highly expressed by MAIT cells at early stages of development in both human and mice (41). The stage-specific increase of H3K27ac at *IKZF1* and *IKZF2* loci, despite limited transcriptional changes, is consistent with epigenetic priming during early MAIT cell development. (42). An earlier study reported that patients lacking Helios have virtually no circulating human MAIT cells, indicating that Helios, along with PLZF, are essential for their development (41).

*BACH2* is highly expressed by immature stage 1 and stage 2 human MAIT cells, and H3K27ac increases at *BACH2* regulatory regions as cells mature to stage 3. *BACH2* acts as a repressor of effector differentiation in naive T cells, and epigenetic profiling of mouse NKT cell development revealed that PLZF bound and repressed *BACH2* (38). Thus, PLZF and *BACH2* expression may represent an important transcriptional switch that occurs as MAIT cells transition through stages 1-3 in the human thymus. In mice, PLZF expression is epigenetically regulated by both repressive (H3K27me3) and active (H3K4me3) histone marks at the *ZBTB16* locus in double-positive thymocytes, creating a poised chromatin state before effector programming (36, 43). The histone methyltransferase EZH2 can co-occupy chromatin with PLZF, and the removal of EZH2 leads to increased binding of PLZF and thus increased expression of PLZF target genes (24). Another study revealed that EZH2 directly methylates lysine 430 on PLZF, leading to its degradation (44). Indeed, the absence of EZH2 in mice leads to much higher frequencies of NKT cells, presumably due to increased binding of PLZF to target genes (43, 44). Contrary to its canonical role in chromatin repression, Koubi et al. reported that EZH2 played a non-canonical role when colocalised with PLZF, associated with active transcription, likely due to the absence of SUZ12 (a core PRC2 component required for H3K27me3 mediated repression) at these binding sites (24). In line with this finding, when we look at regions that contain the PLZF-EZH2 motif (24). We observed a higher H3K27ac signal across MAIT cell differentiation, indicating that PLZF binding at these regions is associated with active chromatin. Collectively, these data suggest a conserved epigenetic mechanism governing PLZF function across subsets of unconventional T cells, including human MAIT cells.

Prior work in humans and mice showed that cytokine production in response to activation is limited to thymic stage 3 human MAIT cells, with further maturation occurring in the blood (8, 10). Consistent with this, we observed enhanced H3K27ac at regulatory regions required for IFNγ expression as human MAIT cells matured in the thymus. In addition, we observed increased H3K27ac and RNA expression at cytokine receptor genes, including *IL18R1* (CD218a), *IL18RAP* (CD218b), *IL2RB* (CD122), *IL7R* (CD127), and *IFNGR1*. These changes suggest that regulatory regions controlling inflammatory signalling pathways become activated during intrathymic MAIT maturation, potentially priming MAIT cells for rapid responses to cytokines such as IL-12 and IL-18 in peripheral tissues (10, 16).

Acetylation was also increased at the *SLAMF6* and *SLAMF7* loci during the development of human MAIT cells. The SLAM–SAP signalling pathway is known to regulate cytotoxicity and cytokine production and is required for innate-like effector differentiation of iNKT and MAIT cells (45–48). SAP-deficient mice showed a severe block of NKT development and a significant reduction of MAIT cells (9, 10, 45, 46, 49). The epigenetic remodelling observed at *SLAMF6* and *SLAMF7* suggests that chromatin level regulation of this signalling pathway is critical for human MAIT cell effector maturation.

Chemokine receptors guide the migration of unconventional T cells within the thymus (10, 50). We detected significant epigenetic changes in chemokine receptor genes during MAIT cell maturation in the thymus. At early stages of development (stage 1 to stage 2), *CCR7* displayed enhanced acetylation and transcript levels, in line with its role in intrathymic migration to the thymic medulla (50, 51). Notably, CCR7-deficient mice have reduced MAIT cells, iNKT cells and γδ T cells (9). Progression of human MAIT cells to stage 3 is accompanied by increased acetylation and transcript levels of *CCR5, CXCR6, CCL4* and *CCL5*, together with upregulation of *S1PR1,* suggesting these molecules may also play important roles during MAIT cell maturation. Studies in mice showed that *CXCR6* is essential for the transition from stage 2 to stage 3 MAIT cells (9, 10), and the absence of *CXCR6* leads to nearly complete loss of mature MAIT cells in peripheral tissues (9, 10). Interestingly, although *CCR9* is highly expressed by immature thymocytes and contributes to the retention of MAIT cells in the thymus (46), we observed increased H3K27ac but decreased transcript levels of *CCR9* during maturation. This is consistent with a developmental switch toward alternative chemokine receptor expression by stage 3 MAIT cells (*CCR5*, *CCR6*, *CXCR6* and *S1PR1*) (52). Together, these epigenetic changes in chemokine receptor loci may regulate the transition of MAIT cells from thymic residency to peripheral migration and tissue homing states. A similar shift in chemokine receptor expression has been reported in Vγ9Vδ2 T cells and NKT cells (13, 17, 35, 53–55), suggesting that chemokine receptor remodelling is a shared feature commonly associated with unconventional T cell development in the thymus.

The pore forming protein in cytotoxic granules, encoded by *PRF1* (perforin), showed increased transcript levels at stage 3 despite reduced H3K27ac. It is currently unclear how these differences in H3K27ac and gene expression translate to perforin expression, but perhaps additional activation signals in the periphery are required for full perforin activity. Increased acetylation and transcript levels at *GZMA* (granzyme A) show that this gene is epigenetically modified and poised to support rapid effector responses. We also identified several genes, including *BCL2*, members of the *GIMAP* family, *CTSW* and *KLF* transcription factors, that were marked by H3K27 acetylation during MAIT cell development. These molecules play key roles in the development and functional maturation of mouse iNKT cells (37), are associated with lymphocyte survival and cytotoxic function (27–29, 37) and thus are also likely to be essential for MAIT cells.

Taken together, our study reveals chromatin remodelling plays a pivotal role in shaping MAIT cell development within the human thymus. Overlap between the transcriptional profiles of MAIT cells, NKT cells, and γδ T cells suggests that conserved epigenetic mechanisms regulate unconventional T cell development. Together, these results suggest that MAIT cells progressively acquire enhancer activity associated with effector function during thymic maturation. Our findings provide new insights into the regulation of genes that control MAIT cell function, and this may accelerate the development of novel approaches to manipulate MAIT cells for therapeutic purposes to treat human disease.

## Materials & Methods

### Study overview

This study focused on the identification of histone modifications (H3K27ac) that potentially influence and regulate the development of MAIT cells. In this investigation, magnetic-activated cell sorting (MACS) enrichment, cell sorting, and cleavage under targets and tagmentation (CUT&Tag) followed by sequencing, were performed to map epigenetic changes of genes, particularly at critical transcription factors in MAIT cells of the postnatal human thymus.

### Sample collection and preparation of a single-cell suspension

Postnatal thymus from children was isolated by a surgeon during cardiac surgery at the Royal Children’s Hospital under ethical approval number (HREC38192), through the Melbourne Children’s Heart Tissue Bank. A total of 7 thymic donors, ranging in age from 7 days to 4 years, were used in this study (full list of donors is provided in supplementary table 11). The extracted thymus samples were cut into tiny pieces and filtered through a 70-μm cell strainer into FACS buffer, which consisted of phosphate-buffered saline (PBS) containing 2% fetal bovine serum (FBS). The resulting mixture was then washed twice before undergoing magnetic bead enrichment.

### Generating MR1 Tetramer

Biotinylated MR1 monomers were obtained from the Corbett McCluskey lab (Dept. Microbiology and Immunology, Peter Doherty Institute, University of Melbourne, Parkville, Australia) and then conjugated to Streptavidin, PE-CF594, step by step, to guarantee that the binding sites of biotin were fully saturated. To achieve the required working concentration, 1.5 mg/mL was diluted with PBS to achieve a final concentration of 0.5 mg/mL. To this solution, Streptavidin-PE-CF594 was added every 5 minutes for a total of 5 additions, with intermittent mixing by gentle pipetting or centrifugation. This final tetramer solution was used for cell staining.

### Isolation of thymic MAIT cells using MACS enrichment

To enrich various developmental stages of MAIT cells, the thymic cell suspensions were initially stained with MR1-5-OP-RU tetramers, and MACS was performed to enrich for MR1-5-OP-RU tetramer-positive cells for the subsequent isolation of stage 1 and stage 2 MAIT cells. To ensure sufficient numbers of CD161^+^ stage 3 MAIT cells, MACS enrichment was performed on CD161-PEVio770^+^ thymic cells and then stained with surface antibodies and MR1-5-OP-RU tetramer as outlined in Supplementary table 13. Magnetic bead enrichment was performed using anti-PE microbeads and LS columns following the manufacturer’s guidelines (Miltenyi Biotec) to isolate sizeable numbers of these cells. Enriched MAIT populations cells were stained with Zombie NIR (viability dye; BioLegend) and the surface antibodies (Supplementary Tables 12 and 13). The staining was conducted on ice for 20 minutes, followed by washing with FACS buffer (PBS enriched with 2% FBS). Subsequently, different stages of MAIT cells were sorted using a BD FACS ARIA Fusion cell sorter and cells were immediately processed for CUT&Tag experiments.

### CUT&Tag experiment

CUT&Tag was performed on approximately 300-2,000 cells for each stage of the MAIT cell development using a modified Epicypher protocol (Henikoff et al.) (56). The reagents and buffers list (Supplementary tables 14-16) was adapted from this EpiCypher CUT&Tag protocol (CUT&Tag Protocol v2.0, revised 7 August 2024. Available at: https://www.epicypher.com/wp-content/uploads/2024/10/cutana-cuttag-protocol.pdf.

Nuclei were isolated using cold NE1 buffer and incubated with activated ConA beads. The cells were incubated overnight at 4°C with the primary antibody against H3K27ac. After washing, a secondary antibody was applied for 30 minutes at room temperature to enhance tagmentation efficiency. The pA-Tn5 transposase loaded with sequencing adapters was added, followed by incubation in tagmentation buffer at 37°C to integrate adapters into fragmented DNA. Reactions were stopped with the SDS Release Buffer, and DNA fragments were purified. PCR was performed to amplify mononucleosome-sized fragments (∼300 bp), which were confirmed using Tape Station traces. Indexed libraries were pooled at equimolar concentrations and sequenced on an Illumina HiSeq 2000 platform.

### Data Analysis

CUT&Tag reads were aligned to the human genome (hg19) using BWA (57), and BAM files were filtered for high-quality reads (mapping quality >15) using SAMtools (58). The Model-based Analysis of ChIP-Seq (MACS2) tool was used to identify H3K27ac-marked regions (peaks, narrow peak settings) (59), and differential peaks were identified with DESeq2 using a significance threshold of P<0.05, fold change (FC) >2, and reads/peak >50 (60). Differential peak data were integrated with published RNA-seq results with the accession number GSE137350 (9) to find any correlations between histone modifications and gene expression. Checking the quality of the CUT&Tag Data was performed using deepTools to generate bigwig files for UCSC Genome Browser visualisation (61). To investigate transcription factor binding motifs (TFBMs) associated with differential peaks, motif enrichment analysis was performed by applying hypergeometric Optimisation of Motif Enrichment (HOMER) (62). This tool recognised enriched motifs, providing insights into potential transcription factors driving gene expression changes during MAIT cell development.

### Pathway Analysis

Enriched pathways and biological processes were identified using the lists of genes associated with differential peaks. For KEGG pathway analysis, gene lists were input into the KEGG Mapper, and significant pathways were visualised with pathway maps. Biological processes were ranked according to their logP values, and the most significant processes were subsequently visualised using bubble plots to elucidate their relevance to MAIT cell differentiation.

## Resource availability

Additional information and requests for resources should be directed to Daniel Pellicci and Boris Novakovic.

## Data and code availability

All H3K27ac CUT&Tag data generated in this study have been deposited in the Gene Expression Omnibus (GEO) under accession number GSE322874. RNA-sequencing data used for integration analyses were previously published (9). Due to file size, the complete integrated H3K27ac and RNA-seq analysis tables, including Supplementary Files 1–3 referenced in this paper, are available via Figshare at: https://doi.org/10.25374/MCRI.31743313 (to be made publicly available upon publication).

## Acknowledgments

We thank Matthew Burton and Eleanor Jones from the Flow Cytometry and Imaging Facility at the Murdoch Children’s Research Institute for their assistance with cell sorting. Marziyeh Taheri is supported by a University of Melbourne post-graduate research scholarship. Louis Perriman is supported by a Don Mulhallen Cancer Research Fellowship from Fiona Elsey Cancer Research Institute. Daniel G Pellicci is supported by a Sylvia & Charles Viertel Fellowship (ViertelSMRF23009). This research was co-funded by the NMHRC ideas grant GNT2030186.

## Author contributions

Conceptualisation, B.N., and D.G.P.; investigation, M.T., B.K., L.P., S.J., C.M., I.E.K., A.T.P., H-F.K., and S.P.B.; formal analysis, M.T., B.K., and B.N.; writing original draft, M.T., L.P., B.N., and D.G.P.; writing – review and editing, M.T., L.P., S.J., C.M., I.E.K., A.T.P., H-F.K., and S.P.B., B.N., and D.G.P.; funding acquisition, S.P.B., B.N., and D.G.P.; supervision, B.N., and D.G.P

