## Supplementary Figures for "Dynamic remodeling of chromatin during human mucosal-associated invariant T cell development"

**Gating strategy**

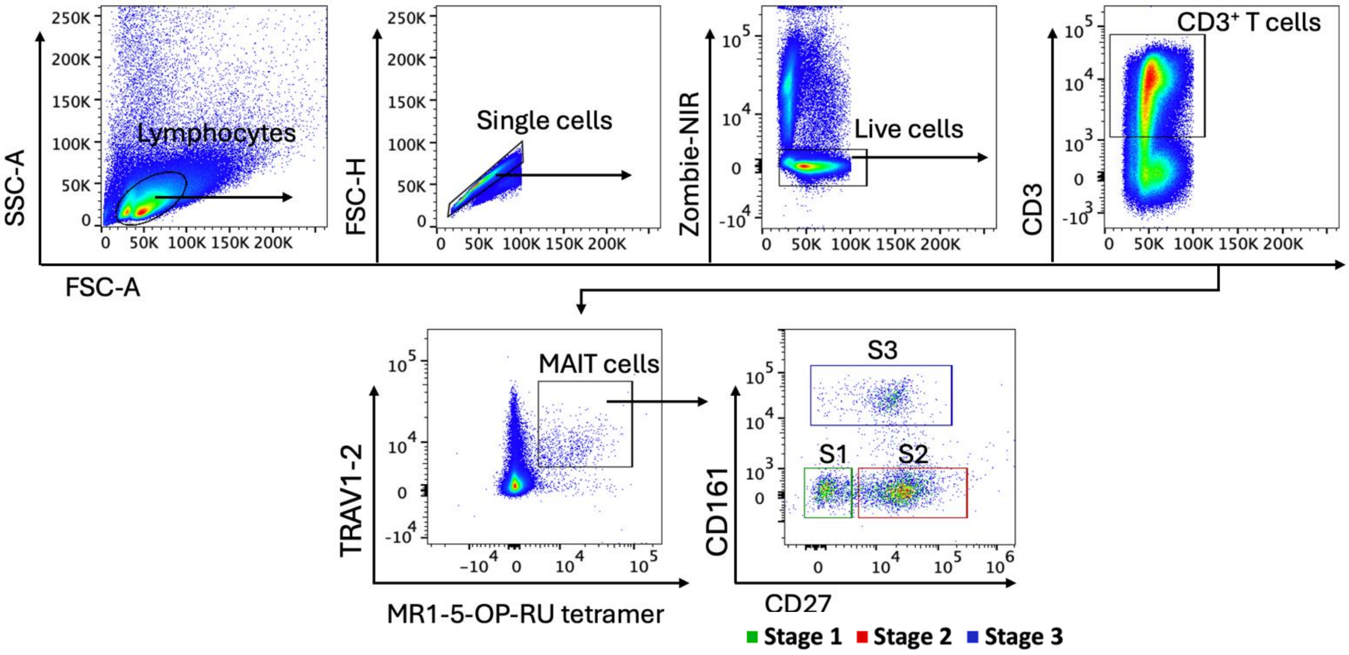


**Supplementary Figure 1. Flow cytometric gating strategy for defining human thymic MAIT cell developmental stages.** Representative flow cytometry plots from a thymus sample of one of the study participants. After gating on lymphocytes, single cells, and live cells, $CD3^{+}$ T cells were selected. From $CD3^{+}$ cells, MAIT cells were identified as MR1-5-OP-RU $tetramer^{+}$ TRAV1-2 (Vα7.2)$^{+}$ cells. Developmental stages of MAIT cells were defined as stage 1 ($CD27^{-}CD161^{-}$) in green, stage 2 ($CD27^{+}CD161^{-}$) in red, and stage 3 ($CD27^{+/-}CD161^{+}$) in blue.

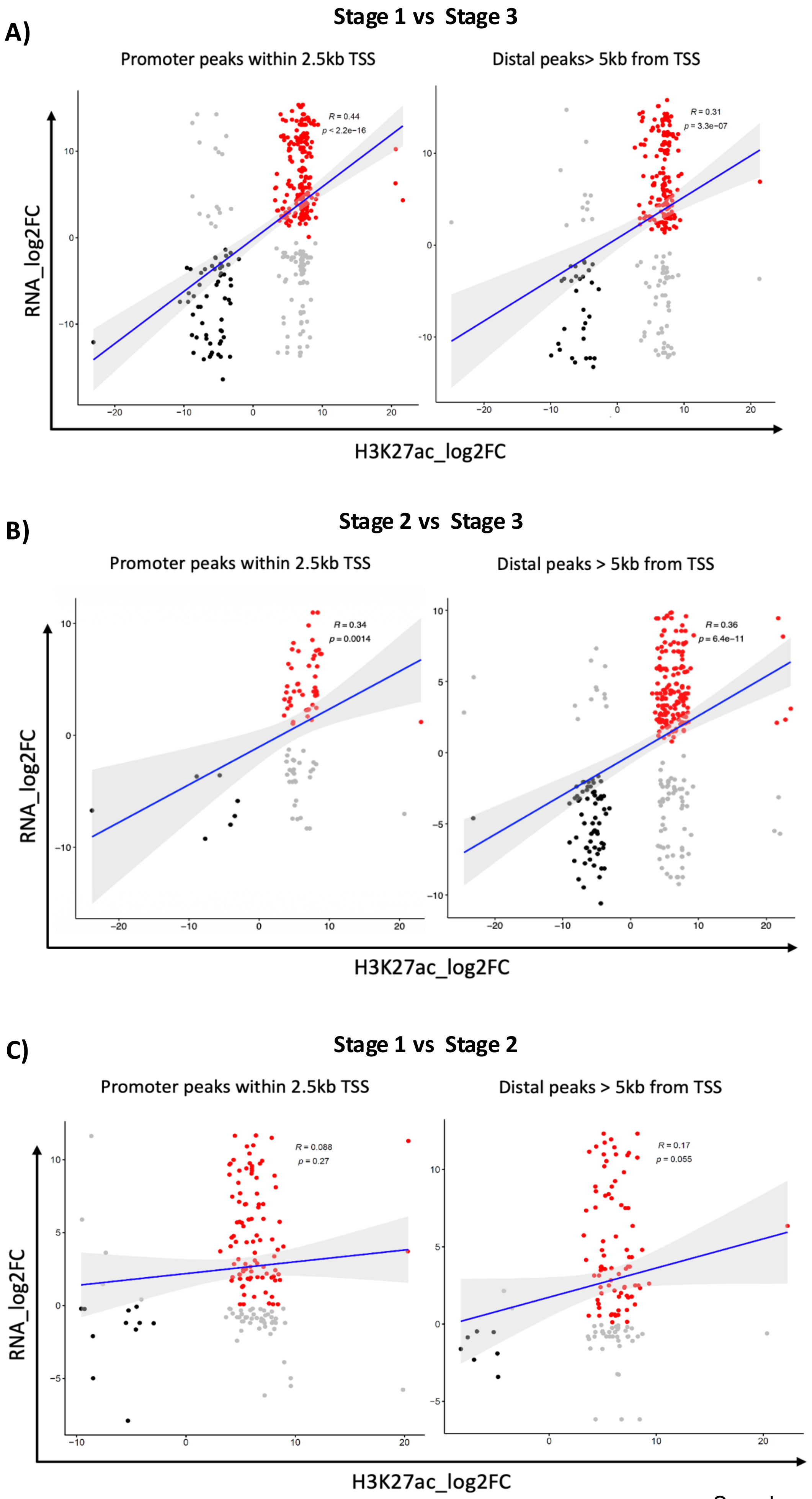


Supplementary figure 2

**Supplementary Figure 2: Correlation of H3K27ac enrichment with gene expression at promoter and distal regions during MAIT cell development.** **(A–C)** Scatter plots show the correlation between changes in H3K27ac signal ($log_2$ fold change, x axis) and dynamic gene expression (CPM $log_2$FC y-axis) at promoter regions (within ± 2.5 kb of the transcription start site (TSS)) and distal regulatory elements (> 5 kb from the TSS) during the transitions from **(A)** stage 1 to stage 3, **(B)** stage 2 to stage 3 and **(C)** stage 1 to stage 2. Each point represents a peak, and only one representative acetylation peak per gene is shown. Red dots indicate genes with increased RNA expression and H3K27ac levels, black dots indicate genes with decreased expression and H3K27ac levels, and grey dots indicate genes that show opposing patterns. Pearson correlation analysis was used to assess the association between H3K27ac and RNA expression. Differential H3K27ac peaks were identified using DESeq2, and RNA expression analysis was performed in R. $P < 0.05$ was considered statistically significant.
