## Supplementary Tables for "Dynamic remodeling of chromatin during human mucosal-associated invariant T cell development"

**Supplementary Table 1.** Genes with decreased H3K27ac and RNA expression in stage 3 compared to stage 1.

**S1 vs S3 Common down (93 genes)**

| Nearest gene | Peak annotation | Nearest gene | Peak annotation | Nearest gene | Peak annotation |
|---|---|---|---|---|---|
| **ACAD9** | Distal | **PLEKHG1** | Distal | **RHOBTB1** | promoter |
| **ADAM10** | Distal | **POU2AF1** | Distal | **SLC16A10** | promoter |
| **AFF2** | Distal | **PPP2R2D** | Distal | **SSFA2** | promoter |
| **APP** | Distal | **RFTN1** | Distal | **ST3GAL6** | promoter |
| **ARPP21** | Distal | **RGS12** | Distal | **VANGL2** | promoter |
| **BCL11A** | Distal | **RORB** | Distal | **ZMYM4** | promoter |
| **BCL2** | Distal | **RPS6KA2** | Distal | **ZNF415** | promoter |
| **NBAS** | Distal | **MTMR1** | Distal | **DSG2** | promoter |
| **PCBP3** | Distal | **PCMTD1** | Distal | **DNTT** | promoter |
| **PCDH9** | Distal | **PCSK5** | Distal | **EGR2** | promoter |
| **PCGF3** | Distal | **PELI2** | Distal | **ELOVL4** | promoter |
| **MAL** | Distal | **MPPED2** | Distal | **PVRIG** | promoter |
| **MTA3** | Distal | **PITPNM2** | Distal | **AXIN2** | promoter |
| **PLCL1** | Distal | **ZNF608** | Distal | **CD109** | promoter |
| **LZTS1** | Distal | **ZNF280D** | Distal | **CD1B** | promoter |
| **MAD1L1** | Distal | **YKT6** | Distal | **EXT1** | promoter |
| **KLF7** | Distal | **WNT5A** | Distal | **FLNB** | promoter |
| **LEF1** | Distal | **UBE2E2** | Distal | **LRP6** | promoter |
| **ITM2A** | Distal | **USP6NL** | Distal | **LRP6** | promoter |
| **CCDC120** | Distal | **SATB1** | Distal | **NDUFA12** | promoter |
| **CCDC6** | Distal | **SCAI** | Distal | **PRPF6** | promoter |
| **CCDC120** | Distal | **SATB1** | Distal | **IQCA1** | Distal |
| **CCDC92** | Distal | **TPCN1** | Distal | **CD1A** | Distal |
| **HHIP** | Distal | **TSNARE1** | Distal | **CD1C** | Distal |
| **ID3** | Distal | **SH2D1A** | Distal | **CECR1** | Distal |
| **SHISA2** | Distal | **SNX29** | Distal | **CEP128** | Distal |
| **SMAD1** | Distal | **SOCS1** | Distal | **DACH1** | Distal |
| **DUSP10** | Distal | **SUSD2** | Distal | **SOX4** | Distal |
| **EPHB6** | Distal | **SYK** | Distal | **SSBP2** | Distal |
| **EVL** | Distal | **TIAM1** | Distal | **TIMP2** | Distal |
| **EXOC4** | Distal | **TOX2** | Distal | **TINAGL1** | Distal |
| **EXOC6** | Distal | **GALNT7** | Distal | **FAM210A** | Distal |

**Supplementary Table 2.** Genes with increased H3K27ac and RNA expression in stage 3 compared to stage 1.

| S1 vs S3 Common up (284 genes) | | | | | | | |
|---|---|---|---|---|---|---|---|
| **Nearest gene** | **Peak annotation** | **Nearest gene** | **Peak annotation** | **Nearest gene** | **Peak annotation** | **Nearest gene** | **Peak annotation** |
| **ACOT7** | Distal | CLN8 | Distal | **IFNG** | Distal | **MYO1F** | Distal |
| **ACTN4** | Distal | **CLPB** | Distal | **IFNGR1** | Distal | **NABP1** | Distal |
| **ADAM19** | Distal | **CMAS** | Distal | **IKZF3** | Distal | **NAGK** | Distal |
| **ADAT2** | Distal | **CNNM4** | Distal | **IL18RAP** | Distal | **NEDD4** | Distal |
| **ADHFE1** | Distal | **CPNE2** | Distal | **IL23R** | Distal | **NEK3** | Distal |
| **ADRB2** | Distal | **CRIP1** | Distal | **IL2RB** | Distal | **NFATC2** | Distal |
| **ADTRP** | Distal | **CSRP1** | Distal | **IL7R** | Distal | **NFKB1** | Distal |
| **AFAP1** | Distal | **CST7** | Distal | **INADL** | Distal | **NOTCH1** | Distal |
| **AGAP7** | Distal | **CTSC** | Distal | **IQGAP2** | Distal | **NTPCR** | Distal |
| **AHNAK** | Distal | **DERL1** | Distal | **IRF1** | Distal | **NUFIP1** | Distal |
| **AIM1** | Distal | **DGKZ** | Distal | **IRF4** | Distal | **PAM** | Distal |
| **AKR1B1** | Distal | **DOCK9** | Distal | **ISG20** | Distal | **PAPOLG** | Distal |
| **ANXA2R** | Distal | **DPP4** | Distal | **ISYNA1** | Distal | **PHTF1** | Distal |
| **APOL1** | Distal | **DTHD1** | Distal | **KIAA0922** | Distal | **PIEZO1** | Distal |
| **ARHGAP18** | Distal | **EIF4E3** | Distal | **KLF3** | Distal | **PIK3AP1** | Distal |
| **ARL6IP6** | Distal | **EMB** | Distal | **KLF6** | Distal | **PLAC8** | Distal |
| **ATG14** | Distal | **EOMES** | Distal | **KLHL2** | Distal | **PPP2R2B** | Distal |
| **ATG16L2** | Distal | **EPHA4** | Distal | **LDLRAP1** | Distal | **PPP2R5C** | Distal |
| **ATP10A** | Distal | **ERN1** | Distal | **LIPC** | Distal | **PREX1** | Distal |
| **B3GNT2** | Distal | **FAM110A** | Distal | **LONRF1** | Distal | **PRKCA** | Distal |
| **BBS9** | Distal | **FAM159A** | Distal | **LPAR6** | Distal | **PRSS35** | Distal |
| **BHLHE40** | Distal | **FAM65B** | Distal | **LPGAT1** | Distal | **PSMA6** | Distal |
| **BIRC3** | Distal | **FEM1B** | Distal | **LPIN1** | Distal | **PTCH1** | Distal |
| **BLK** | Distal | **FLT4** | Distal | **LRIG1** | Distal | **PTGDR** | Distal |
| **BTN2A1** | Distal | **FOSB** | Distal | **LRP8** | Distal | **PTPN22** | Distal |
| **C10orf54** | Distal | **FOSL2** | Distal | **LTBP4** | Distal | **PTPRJ** | Distal |
| **C11orf21** | Distal | **FOXJ2** | Distal | **LYAR** | Distal | **PWP2** | Distal |
| **C6orf57** | Distal | **FOXO1** | Distal | **MAF** | Distal | **RAB27A** | Distal |
| **CACNA2D1** | Distal | **FUT11** | Distal | **MAN1C1** | Distal | **RAB30** | Distal |
| **CAMSAP1** | Distal | **FXN** | Distal | **MAP3K1** | Distal | **RALA** | Distal |
| **CAPN2** | Distal | **GAB3** | Distal | **MAP3K5** | Distal | **RGS3** | Distal |
| **CBLB** | Distal | **GALC** | Distal | **MAPKAPK3** | Distal | **RHOB** | Distal |
| **CBX7** | Distal | **GBP5** | Distal | **MAR08** | Distal | **RNF6** | Distal |
| **CCL5** | Distal | **GIMAP5** | Distal | **MATN2** | Distal | **RORA** | Distal |

| | | | | | | | |
|---|---|---|---|---|---|---|---|
| **CCND2** | Distal | **GIMAP7** | Distal | **MBOAT1** | Distal | **RUNX3** | Distal |
| **CD244** | Distal | **GLUD1** | Distal | **MBP** | Distal | **RXRA** | Distal |
| **CD74** | Distal | **GPR183** | Distal | **METAP1D** | Distal | **S100B** | Distal |
| **CD97** | Distal | **GPR68** | Distal | **MPP7** | Distal | **S1PR1** | Distal |
| **CDC14A** | Distal | **HERC6** | Distal | **MX1** | Distal | **LAX1** | promoter |
| **CEBPD** | Distal | **HM13** | Distal | **SAMD3** | Distal | **TC2N** | promoter |
| **CHD7** | Distal | **HOPX** | Distal | **WDR75** | Distal | **TMEM192** | promoter |
| **CLIC5** | Distal | **IFI44** | Distal | **LRRC8B** | promoter | **TNFSF14** | promoter |
| **SEL1L3** | Distal | **MX2** | Distal | **MARS2** | promoter | **TPI1** | promoter |
| **SEMA4C** | Distal | **MYBL1** | Distal | **MBTPS2** | promoter | **ULK3** | promoter |
| **SHISA5** | Distal | **MYC** | Distal | **MDFIC** | promoter | **VAMP8** | promoter |
| **SHQ1** | Distal | **SAV1** | Distal | **MT1X** | promoter | **ZC3HAV1L** | promoter |
| **SIPA1L2** | Distal | **SCAMP2** | Distal | **MT2A** | promoter | **ZNF561** | promoter |
| **SKI** | Distal | **SDAD1** | Distal | **MTIF2** | promoter | **ZNF674** | promoter |
| **SLAMF6** | Distal | **ZC3H7A** | Distal | **NFKBIZ** | promoter | **IL18R1** | promoter |
| **SLAMF7** | Distal | **ZEB2** | Distal | **PCNXL2** | promoter | **TBK1** | promoter |
| **SLC15A4** | Distal | **ZFP64** | Distal | **ACOT9** | promoter | **LAG3** | promoter |
| **SLC22A5** | Distal | **ZNF680** | Distal | **ANXA5** | promoter | **TBC1D10B** | promoter |
| **SMAD7** | Distal | **ZNF831** | Distal | **ARMC2** | promoter | **IL15RA** | promoter |
| **SNTB1** | Distal | **ZYX** | Distal | **C4orf32** | promoter | **SMURF1** | promoter |
| **SORL1** | Distal | **WIPF1** | Distal | **CCL4** | promoter | **SNRPA** | promoter |
| **SPATS2L** | Distal | **XPNPEP1** | Distal | **PHACTR2** | promoter | **SNX25** | promoter |
| **SRGN** | Distal | **YPEL4** | Distal | **PIM2** | promoter | **SP140** | promoter |
| **SRPRB** | Distal | **ZBP1** | Distal | **PLEK** | promoter | **SPOPL** | promoter |
| **SSBP4** | Distal | **ZBTB16** | Distal | **PPAP2A** | promoter | **SPRY2** | promoter |
| **STAT4** | Distal | **CCR5** | promoter | **PRDM15** | promoter | **SRBD1** | promoter |
| **STK10** | Distal | **CLSTN3** | promoter | **PRR7** | promoter | **STOM** | promoter |
| **SVIP** | Distal | **CRTAP** | promoter | **RAB7L1** | promoter | **SLC3A2** | promoter |
| **SYTL2** | Distal | **CTSW** | promoter | **RNF135** | promoter | **SLC4A4** | promoter |
| **TGFBR3** | Distal | **CUL5** | promoter | **S1PR4** | promoter | **TRAPPC2** | Distal |
| **TGIF1** | Distal | **CXCR6** | promoter | **SAMHD1** | promoter | **DNAL1** | promoter |
| **TIFA** | Distal | **CYBA** | promoter | **SLC25A12** | promoter | **DVL3** | promoter |
| **TMEM127** | Distal | **DHX57** | promoter | **SLC36A1** | promoter | **EDC3** | promoter |
| **EMP3** | promoter | **FNDC3B** | promoter | **IFIT1** | promoter | **TSPAN5** | Distal |
| **FAM46A** | promoter | **GIMAP6** | promoter | **UTP18** | Distal | **TXNIP** | Distal |
| **FAS** | promoter | **GNPDA1** | promoter | **VDAC1** | Distal | **TSC22D3** | Distal |
| **TRIM14** | Distal | **TNFRSF1B** | Distal | **TMEM200A** | Distal | **TNFAIP3** | Distal |
| **TRIM69** | Distal | | | | | | |

**Supplementary Table 3.** Genes with mixed H3K27ac and RNA expression changes between Stage 1 and Stage 3.

| S1 vs S3 Mixed direction (172 genes) | | | | | | | |
|---|---|---|---|---|---|---|---|
| **Nearest gene** | **Peak annotation** | **Nearest gene** | **Peak annotation** | **Nearest gene** | **Peak annotation** | **Nearest gene** | **Peak annotation** |
| **PPP2R2D** | Distal | **SOCS1** | Distal | **AKAP11** | Distal | **RPS14** | Distal |
| **SIRPG** | Distal | **CDK18** | Distal | **BRD1** | Distal | **KLF6** | Distal |
| **LANCL2** | Distal | **FAM210A** | Distal | **ITM2A** | Distal | **STOM** | Distal |
| **GRK5** | Distal | **EVL** | Distal | **CD9** | Distal | **USO1** | Distal |
| **ANKDD1A** | Distal | **CCDC6** | Distal | **CMTM6** | Distal | **FTO** | Distal |
| **TIAM1** | Distal | **TMEM185B** | Distal | **LZTS1** | Distal | **SLC1A1** | Distal |
| **HADH** | Distal | **CDK6** | Distal | **DLD** | Distal | **TMC6** | Distal |
| **RHOH** | Distal | **HIVEP3** | Distal | **APP** | Distal | **CHN2** | Distal |
| **LGMN** | Distal | **ABCD1** | Distal | **COPS8** | Distal | **TMEM200A** | Distal |
| **IQCA1** | Distal | **LIG3** | Distal | **LYRM4** | Distal | **BCL2** | Distal |
| **RFTN1** | Distal | **GCNT4** | Distal | **SOX4** | Distal | **EXOC6** | Distal |
| **YKT6** | Distal | **THEMIS** | Distal | **CSRNP1** | Distal | **MEF2D** | Distal |
| **CHST2** | Distal | **RGS12** | Distal | **SIRPB1** | Distal | **ANAPC7** | Distal |
| **ASB7** | Distal | **ENC1** | Distal | **DCK** | Distal | **MTMR1** | Distal |
| **SUPT3H** | Distal | **HDAC7** | Distal | **ALDH7A1** | Distal | **SH2D1A** | Distal |
| **CHI3L2** | Distal | **ZEB1** | Distal | **MKL2** | Distal | **CD79A** | Distal |
| **NINJ2** | Distal | **BCL11A** | Distal | **ADAM10** | Distal | **PAWR** | Distal |
| **KATNB1** | Distal | **VANGL2** | Distal | **DPCD** | Distal | **MFI2** | Distal |
| **STRBP** | Distal | **KLHL3** | Distal | **RUFY3** | Distal | **NT5E** | Distal |
| **IGF2R** | Distal | **GFOD1** | Distal | **LEF1** | Distal | **DDX60** | Distal |
| **FLI1** | Distal | **SNX29** | Distal | **RGCC** | Distal | **ACYP2** | Distal |
| **ZNF608** | Distal | **GPR125** | Distal | **SHISA2** | Distal | **RAG1** | Distal |
| **FLNB** | Distal | **ELF4** | Distal | **GRAP2** | Distal | **CST7** | Distal |
| **PGS1** | Distal | **TBC1D4** | Distal | **ZNF92** | Distal | **DHX37** | promoter |
| **CD226** | Distal | **PELI1** | Distal | **ZFYVE21** | Distal | **TMSB15B** | promoter |
| **PET112** | Distal | **CECR1** | Distal | **FOXO1** | Distal | **MRPS11** | promoter |
| **MAD1L1** | Distal | **KATNAL1** | Distal | **LPIN1** | Distal | **PVRIG** | promoter |
| **RTN4R** | Distal | **CCR9** | Distal | **ARHGAP18** | Distal | **SIT1** | promoter |
| **RNF24** | Distal | **EYA3** | Distal | **AGPAT3** | Distal | **RNASE6** | promoter |
| **TCF7** | Distal | **ZNF236** | Distal | **ADAT2** | Distal | **CPEB2** | promoter |
| **EGR2** | Distal | **DPPA4** | Distal | **CLN8** | Distal | **DRG2** | promoter |
| **TMCO3** | Distal | **ACPL2** | Distal | **NTPCR** | Distal | **NSD1** | promoter |
| **NBPF9** | Distal | **SLC44A1** | Distal | **SAMD3** | Distal | **NUCB2** | promoter |
| **IMPG2** | Distal | **SFSWAP** | Distal | **CLEC2B** | Distal | **TMEM159** | promoter |
| **RHOU** | Distal | **SUSD2** | Distal | **PRF1** | Distal | **TRAF7** | promoter |

| | | | | | | | |
|---|---|---|---|---|---|---|---|
| **VILL** | Distal | **TOX2** | Distal | **CD101** | Distal | **TCEAL8** | promoter |
| **KDM2B** | Distal | **SEC14L1** | Distal | **TSC22D3** | Distal | **USP40** | promoter |
| **BACH2** | Distal | **CEBPZ** | Distal | **RAPGEF2** | Distal | **RDH14** | promoter |
| **SRD5A3** | promoter | **FAM84B** | promoter | **ZBTB37** | promoter | **GIMAP7** | promoter |
| **KIAA1009** | promoter | **RFX3** | promoter | **PIK3C3** | promoter | **CAMSAP2** | promoter |
| **IFITM1** | promoter | **LAX1** | promoter | **NXPE3** | promoter | **GLTP** | promoter |
| **C17orf75** | promoter | **NUP133** | Promoter | **USP38** | Promoter | **SERPINF1** | promoter |
| **ZSWIM6** | promoter | **ZNF407** | promoter | **FAN1** | promoter | | |

**Supplementary Table 4.** Genes with decreased H3K27ac and RNA expression in stage 2 compared to stage 1.

| **S1 vs S2 Common down (42 genes)** | | | |
|---|---|---|---|
| **Nearest gene** | **Peak annotation** | **Nearest gene** | **Peak annotation** |
| **B4GALT1** | Distal | **SPRY1** | Distal |
| **CASP8** | Distal | **TIGD2** | Distal |
| **CEP128** | Distal | **TPD52** | Distal |
| **COPS8** | Distal | **TPK1** | Distal |
| **CYB561D1** | Distal | **USP7** | Distal |
| **DDX11** | Distal | **CCDC28B** | Promoter |
| **DPYSL2** | Distal | **DAPK1** | Promoter |
| **ELF4** | Distal | **DDB2** | Promoter |
| **FBXL14** | Distal | **DRG1** | Promoter |
| **HENMT1** | Distal | **FAM73A** | Promoter |
| **INSIG1** | Distal | **FOXN2** | Promoter |
| **KLHL25** | Distal | **KLHL24** | Promoter |
| **LINGO4** | Distal | **MCMBP** | Promoter |
| **LMO4** | Distal | **NFXL1** | Promoter |
| **MGMT** | Distal | **PDIA6** | Promoter |
| **NEDD9** | Distal | **PIGO** | Promoter |
| **PITPNM2** | Distal | **RBL1** | Promoter |
| **PTPRM** | Distal | **RCSD1** | Promoter |
| **RORB** | Distal | **SPRED2** | Promoter |
| **SGMS1** | Distal | **TTK** | Promoter |
| **SLC35E3** | Distal | **ZNRF1** | Promoter |

**Supplementary Table 5.** Genes with increased H3K27ac and RNA expression in stage 2 compared to stage 1.

**S1 vs S2 Common up (138 genes)**

| Nearest gene | Peak annotation | Nearest gene | Peak annotation | Nearest gene | Peak annotation | Nearest gene | Peak annotation |
|---|---|---|---|---|---|---|---|
| **AGAP7** | Distal | **EIF3J** | Distal | **RHOB** | Distal | **AXIN1** | Promoter |
| **AIM1** | Distal | **FAM65B** | Distal | **RUNX3** | Distal | **B3GNTL1** | Promoter |
| **AMPD3** | Distal | **FARS2** | Distal | **S100B** | Distal | **C12orf4** | Promoter |
| **APOL1** | Distal | **FEM1B** | Distal | **S1PR1** | Distal | **CLSTN3** | Promoter |
| **ARL6IP6** | Distal | **FLT4** | Distal | **SCAMP2** | Distal | **CTSW** | Promoter |
| **ATG14** | Distal | **FOXO1** | Distal | **SEMA4C** | Distal | **DNAJC16** | Promoter |
| **ATP10A** | Distal | **FXN** | Distal | **SGIP1** | Distal | **EIF2B1** | Promoter |
| **B3GNT2** | Distal | **GLUD1** | Distal | **SHQ1** | Distal | **FAM46A** | Promoter |
| **BBS9** | Distal | **GNPDA1** | Distal | **SLAMF6** | Distal | **FAS** | Promoter |
| **BCL11A** | Distal | **GYPC** | Distal | **SLC15A4** | Distal | **GIMAP6** | Promoter |
| **BTLA** | Distal | **INADL** | Distal | **SLC22A5** | Distal | **GPSM2** | Promoter |
| **BTN2A1** | Distal | **ISYNA1** | Distal | **SLC25A33** | Distal | **IFI44** | Promoter |
| **C11orf21** | Distal | **KCTD5** | Distal | **SLCO4A1** | Distal | **INTS2** | Promoter |
| **C6orf57** | Distal | **LDLRAP1** | Distal | **SORL1** | Distal | **LRRC8B** | Promoter |
| **CARD11** | Distal | **LHPP** | Distal | **SRGN** | Distal | **PHACTR2** | Promoter |
| **CCDC64** | Distal | **LIPC** | Distal | **SSBP4** | Distal | **PIM2** | Promoter |
| **CCDC88C** | Distal | **LPAR6** | Distal | **STAT4** | Distal | **PLRG1** | Promoter |
| **CCND2** | Distal | **LRP8** | Distal | **STK10** | Distal | **PPAP2A** | Promoter |
| **CCR7** | Distal | **LTBP4** | Distal | **SYTL2** | Distal | **PPAPDC1B** | Promoter |
| **CD74** | Distal | **MAML3** | Distal | **TC2N** | Distal | **PRDM15** | Promoter |
| **CD97** | Distal | **MAN1C1** | Distal | **TMEM127** | Distal | **PRR7** | Promoter |
| **CLPB** | Distal | **MAP3K1** | Distal | **TMEM200A** | Distal | **RNF135** | Promoter |
| **CLYBL** | Distal | **MAPKAPK3** | Distal | **TNFAIP3** | Distal | **SAMHD1** | Promoter |
| **CRIP1** | Distal | **MBP** | Distal | **TNFRSF1B** | Distal | **SLC25A12** | Promoter |
| **CSRP1** | Distal | **MPP7** | Distal | **TNFSF10** | Distal | **SLC25A37** | Promoter |
| **CST7** | Distal | **NFATC1** | Distal | **TRIM14** | Distal | **SLC36A1** | Promoter |
| **CTSC** | Distal | **NFATC2** | Distal | **UTP18** | Distal | **SPRY2** | Promoter |
| **CYTH4** | Distal | **NUFIP1** | Distal | **ZBTB16** | Distal | **STOM** | Promoter |
| **DAP** | Distal | **PDP1** | Distal | **ZNF518B** | Distal | **TBC1D10B** | Promoter |
| **DAPK2** | Distal | **PHB2** | Distal | **ZNF680** | Distal | **TNFSF12** | Promoter |
| **DBH** | Distal | **PHTF1** | Distal | **ZNF831** | Distal | **ULK3** | Promoter |
| **DERL1** | Distal | **PLEKHA2** | Distal | **ZSWIM4** | Distal | **VAMP8** | Promoter |
| **DIP2A** | Distal | **PPP2R2B** | Distal | **ZYX** | Distal | **YPEL4** | Promoter |
| **DPM2** | Distal | **PWP2** | Distal | **AHNAK** | Promoter | **ARL4A** | Promoter |
| **EDAR** | Distal | **REL** | Distal | | | | |

**Supplementary Table 6.** Genes with mixed H3K27ac and RNA expression changes between Stage 1 and Stage 2.

| S1 vs S2 Mixed direction (117 genes) | | | | | |
|---|---|---|---|---|---|
| **Nearest gene** | **Peak annotation** | **Nearest gene** | **Peak annotation** | **Nearest gene** | **Peak annotation** |
| **ADAM19** | Distal | **KATNAL1** | Distal | **STX18** | Distal |
| **ADRBK2** | Distal | **KLHL25** | Distal | **SYTL2** | Distal |
| **AKR1B1** | Distal | **LITAF** | Distal | **TIGIT** | Distal |
| **ANAPC7** | Distal | **LMNB1** | Distal | **TNFAIP3** | Distal |
| **ANKDD1A** | Distal | **LRP2BP** | Distal | **TNFSF10** | Distal |
| **ASB6** | Distal | **MAP2K3** | Distal | **TYW5** | Distal |
| **B3GNT2** | Distal | **MAP3K1** | Distal | **ZFYVE28** | Distal |
| **B4GALT1** | Distal | **MAP3K5** | Distal | **ZHX1** | Distal |
| **BCL11A** | Distal | **MBP** | Distal | **ZNF704** | Distal |
| **BOD1L1** | Distal | **MFI2** | Distal | **ZYX** | Distal |
| **CACNA2D1** | Distal | **MGMT** | Distal | **JAZF1** | Distal |
| **CACNA2D2** | Distal | **MORN3** | Distal | **STK10** | Distal |
| **CARD11** | Distal | **MTSS1L** | Distal | **ADAM22** | Promoter |
| **CASP8** | Distal | **MYOM2** | Distal | **ANKRD52** | Promoter |
| **CCND2** | Distal | **NAB1** | Distal | **C17orf75** | Promoter |
| **CDK18** | Distal | **NARG2** | Distal | **C3orf52** | Promoter |
| **CDK6** | Distal | **NEDD9** | Distal | **COMTD1** | Promoter |
| **CHN2** | Distal | **PELO** | Distal | **DCUN1D2** | Promoter |
| **COPS8** | Distal | **PHTF1** | Distal | **FAM65B** | Promoter |
| **CTSC** | Distal | **PLEKHA2** | Distal | **GCNT2** | Promoter |
| **CYTH4** | Distal | **PLXND1** | Distal | **GLUD1** | Promoter |
| **DAP** | Distal | **PRPS1** | Distal | **IDI1** | Promoter |
| **DUSP10** | Distal | **PRR5L** | Distal | **IRF2** | Promoter |
| **DUSP16** | Distal | **RANBP10** | Distal | **LMO4** | Promoter |
| **DUSP7** | Distal | **RAPGEF2** | Distal | **MAML3** | Promoter |
| **ELF4** | Distal | **RASGRF2** | Distal | **MTHFD1L** | Promoter |
| **ELK3** | Distal | **RCSD1** | Distal | **NDUFA10** | Promoter |
| **EZH2** | Distal | **RORC** | Distal | **NFIL3** | Promoter |
| **FARS2** | Distal | **RUNX3** | Distal | **PDIA6** | Promoter |
| **FLT4** | Distal | **S1PR1** | Distal | **PLEKHO1** | Promoter |
| **FOXN2** | Distal | **SGIP1** | Distal | **PUS1** | Promoter |
| **FOXO1** | Distal | **SGMS1** | Distal | **SLC35B1** | Promoter |
| **GTDC1** | Distal | **SLC25A24** | Distal | **SLC36A4** | Promoter |
| **GYPC** | Distal | **SLC39A11** | Distal | **TIPARP** | Promoter |
| **HADH** | Distal | **SMAP2** | Distal | **TPK1** | Promoter |

| | | | | | |
|---|---|---|---|---|---|
| **HIVEP2** | Distal | **SNUPN** | Distal | **TRIM14** | Promoter |
| **HVCN1** | Distal | **SORL1** | Distal | **TSC22D3** | Promoter |
| **ISCU** | Distal | **SPTBN1** | Distal | **YAF2** | Promoter |
| **ZRANB3** | Promoter | **ACBD4** | Promoter | **ACBD5** | Promoter |
| **ZRANB3** | Promoter | | | | |

**Supplementary Table 7.** Genes with decreased H3K27ac and RNA expression in stage 3 compared to stage 2.

| **S2 vs S3 Common down (53 genes)** | | | | | | | |
|---|---|---|---|---|---|---|---|
| **Nearest gene** | **Peak annotation** | **Nearest gene** | **Peak annotation** | **Nearest gene** | **Peak annotation** | **Nearest gene** | **Peak annotation** |
| **ACPL2** | Distal | **PITPNM2** | Distal | **PCSK5** | Distal | **ZNF415** | Promoter |
| **AFAP1L1** | Distal | **PLXDC1** | Distal | **PCSK5** | Distal | **ZNF415** | Promoter |
| **AFF2** | Distal | **PPP2R2D** | Distal | **NDNF** | Distal | **SLC16A10** | Promoter |
| **AXIN2** | Distal | **PRKCE** | Distal | **NDNF** | Distal | **SLC16A10** | Promoter |
| **BCL11A** | Distal | **PRL** | Distal | **NDNF** | Distal | **SLC16A10** | Promoter |
| **CCDC6** | Distal | **RAD54L2** | Distal | **NBAS** | Distal | **MBOAT2** | Promoter |
| **CD1A** | Distal | **RFTN1** | Distal | **MAML3** | Distal | **LRP6** | Promoter |
| **CD1C** | Distal | **SATB1** | Distal | **MAL** | Distal | **CD1B** | Promoter |
| **CHST2** | Distal | **SHISA2** | Distal | **MAD1L1** | Distal | **ZNF608** | Distal |
| **CLDN1** | Distal | **SOX4** | Distal | **LZTS1** | Distal | **ZNF280D** | Distal |
| **DACH1** | Distal | **SSBP2** | Distal | **LRRN3** | Distal | **WNT5A** | Distal |
| **EPHB6** | Distal | **SULT1B1** | Distal | **LRRC16A** | Distal | **TPCN1** | Distal |
| **EXOC6** | Distal | **SUPT3H** | Distal | **LEF1** | Distal | **TOX2** | Distal |
| **IQCA1** | Distal | **TIAM1** | Distal | **LCP2** | Distal | **TIMP2** | Distal |
| **PELI2** | Distal | | | | | | |

**Supplementary Table 8.** Genes with increased H3K27ac and RNA expression in stage 3 compared to stage 2.

| **S2 vs S3 Common up (150 genes)** | | | | | | | |
|---|---|---|---|---|---|---|---|
| **Nearest gene** | **Peak annotation** | **Nearest gene** | **Peak annotation** | **Nearest gene** | **Peak annotation** | **Nearest gene** | **Peak annotation** |
| **ACBD5** | Distal | **FOSB** | Distal | **ODC1** | Distal | **ZEB2** | Distal |
| **ACTN4** | Distal | **GAB3** | Distal | **PHPT1** | Distal | **ZNF429** | Distal |
| **ADAM12** | Distal | **GALC** | Distal | **PIK3AP1** | Distal | **RF1** | Distal |
| **ADHFE1** | Distal | **GBP5** | Distal | **PLAC8** | Distal | **ACTR6** | Promoter |
| **ADRB2** | Distal | **GCNT2** | Distal | **PPP2R5C** | Distal | **ANXA2** | Promoter |
| **AHNAK** | Distal | **GIMAP7** | Distal | **PREX1** | Distal | **C11orf24** | Promoter |
| **ALDH18A1** | Distal | **GOLGA8A** | Distal | **PRR5** | Distal | **CCL4** | Promoter |
| **ANKH** | Distal | **GPR183** | Distal | **PRSS35** | Distal | **CCR5** | Promoter |

| | | | | | | | |
|---|---|---|---|---|---|---|---|
| **APOBEC3G** | Distal | **GPR68** | Distal | **PTGDR** | Distal | **CDC27** | Promoter |
| **ARMC2** | Distal | **GTF3A** | Distal | **PTPN22** | Distal | **CLSTN3** | Promoter |
| **B4GALT1** | Distal | **GZMA** | Distal | **PTPRM** | Distal | **CXCR6** | Promoter |
| **BHLHE40** | Distal | **HEG1** | Distal | **RAB27A** | Distal | **DDB2** | Promoter |
| **BLK** | Distal | **HENMT1** | Distal | **RAG1** | Distal | **DUS4L** | Promoter |
| **C1GALT1** | Distal | **HOPX** | Distal | **RGS3** | Distal | **FAM73A** | Promoter |
| **CASP8** | Distal | **IFNGR1** | Distal | **RRAS2** | Distal | **FPGS** | Promoter |
| **CBLB** | Distal | **IL18RAP** | Distal | **RUNX3** | Distal | **GABARAPL1** | Promoter |
| **CCL5** | Distal | **IL23R** | Distal | **SERPINB1** | Distal | **GPX7** | Promoter |
| **CDC14A** | Distal | **IL7R** | Distal | **SESN1** | Distal | **GYG1** | Promoter |
| **CDC42EP3** | Distal | **INSIG1** | Distal | **SETD7** | Distal | **HRAS** | Promoter |
| **CEBPD** | Distal | **JUNB** | Distal | **SGMS1** | Distal | **IFT57** | Promoter |
| **CHD7** | Distal | **KCNQ3** | Distal | **SLC10A7** | Distal | **IL18R1** | Promoter |
| **CLN8** | Distal | **KLF3** | Distal | **SMAD7** | Distal | **LAG3** | Promoter |
| **CST7** | Distal | **KLF6** | Distal | **SNTB1** | Distal | **ME1** | Promoter |
| **CXCR4** | Distal | **KLHDC4** | Distal | **SNX25** | Distal | **MT1X** | Promoter |
| **DDX43** | Distal | **KLHL25** | Distal | **SREBF1** | Distal | **MT2A** | Promoter |
| **DPP4** | Distal | **KLRB1** | Distal | **SREK1IP1** | Distal | **NDUFAF5** | Promoter |
| **DUSP7** | Distal | **LINGO4** | Distal | **STAT4** | Distal | **PDIA6** | Promoter |
| **EIF4E3** | Distal | **LRIG1** | Distal | **TGFBR3** | Distal | **PLEK** | Promoter |
| **ELK3** | Distal | **LTBP2** | Distal | **TGIF1** | Distal | **PRDM1** | Promoter |
| **EOMES** | Distal | **MAF** | Distal | **TMEM43** | Distal | **S100A6** | Promoter |
| **EPHA4** | Distal | **MBOAT1** | Distal | **TMPRSS3** | Distal | **SEL1L3** | Promoter |
| **ERN1** | Distal | **MICALCL** | Distal | **TPK1** | Distal | **SLC9B2** | Promoter |
| **FAM110A** | Distal | **MTSS1L** | Distal | **TRAF3IP2** | Distal | **SRSF4** | Promoter |
| **FAM3C** | Distal | **MYC** | Distal | **TUBD1** | Distal | **WRNIP1** | Promoter |
| **FAM46C** | Distal | **MYO1F** | Distal | **USP13** | Distal | **ZNF668** | Promoter |
| **FASLG** | Distal | **NBEAL2** | Distal | **WIPF1** | Distal | **ACOT9** | Promoter |
| **FLNA** | Distal | **NCALD** | Distal | **WWC3** | Distal | | |
| **WDR4** | Distal | **VDAC1** | Distal | **ZBP1** | Distal | | |

**Supplementary Table 9.** Genes with mixed H3K27ac and RNA expression changes between Stage 2 and Stage 3.

| **S2 vs S3 Mixed direction (87)** | | | | | |
|---|---|---|---|---|---|
| **Nearest gene** | **Peak annotation** | **Nearest gene** | **Peak annotation** | **Nearest gene** | **Peak annotation** |
| **ABLIM1** | Distal | **L3MBTL3** | Distal | **UBIAD1** | Distal |
| **ADRBK2** | Distal | **LCLAT1** | Distal | **USP6NL** | Distal |
| **AKAP11** | Distal | **LCP2** | Distal | **VANGL2** | Distal |
| **ALDH5A1** | Distal | **LGMN** | Distal | **VRK2** | Distal |
| **ARHGAP18** | Distal | **LOH12CR1** | Distal | **ZNF608** | Distal |

| | | | | | |
|---|---|---|---|---|---|
| **B4GALT3** | Distal | **LZTS1** | Distal | **ZNF704** | Distal |
| **BCL11A** | Distal | **MAD1L1** | Distal | **ADNP2** | Promoter |
| **BCL2** | Distal | **MCTP1** | Distal | **APBB1** | Promoter |
| **CCNG2** | Distal | **MTSS1L** | Distal | **CAPRIN1** | Promoter |
| **CECR1** | Distal | **NBAS** | Distal | **CD302** | Promoter |
| **CLN8** | Distal | **NBPF9** | Distal | **CRBN** | Promoter |
| **COA5** | Distal | **PELI1** | Distal | **CSRNP1** | Promoter |
| **COL6A3** | Distal | **PET112** | Distal | **DHX37** | Promoter |
| **CST7** | Distal | **PLAG1** | Distal | **EXT1** | Promoter |
| **DGCR8** | Distal | **RNF4** | Distal | **IFT80** | Promoter |
| **DLD** | Distal | **SGIP1** | Distal | **IFT88** | Promoter |
| **DPM2** | Distal | **SHISA2** | Distal | **JUP** | Promoter |
| **ENC1** | Distal | **SIRPB1** | Distal | **LRRC16A** | Promoter |
| **FLNB** | Distal | **SIRPG** | Distal | **MMS22L** | Promoter |
| **FRY** | Distal | **SIT1** | Distal | **NMRAL1** | Promoter |
| **GRAMD3** | Distal | **SLC39A11** | Distal | **PARP6** | Promoter |
| **GRK5** | Distal | **SLCO4A1** | Distal | **PATL1** | Promoter |
| **HHAT** | Distal | **SPTBN1** | Distal | **RIMKLB** | Promoter |
| **HIVEP3** | Distal | **STARD5** | Distal | **SEC23IP** | Promoter |
| **IGF1R** | Distal | **STOM** | Distal | **SGTB** | Promoter |
| **IGF2R** | Distal | **SUPT3H** | Distal | **STAT5B** | Promoter |
| **KATNAL1** | Distal | **TMSB15B** | Distal | **TEP1** | Promoter |
| **KIAA1324L** | Distal | **TRAF3IP2** | Distal | **TTYH3** | Promoter |
| **KLF6** | Distal | **UBAP1** | Distal | **YLPM1** | Promoter |

**Supplementary Table 10.** Transcription factors with increased or decreased H3K27ac and RNA expression (Maturity and Immaturity).

| **Transcription factors (148)** | | | |
|---|---|---|---|
| **Gene_Name** | **Family** | **Group** | **Dynamic** |
| **PHTF1** | PHTF | Maturity_TF | SHARP |
| **KLF3** | C2H2 Kruepel | Maturity_TF | SHARP |
| **FOXJ2** | Forkhead | Maturity_TF | SHARP |
| **SMAD7** | MAD | Maturity_TF | SLOWER |
| **NOTCH1** | NOTCH | Maturity_TF | SHARP |
| **BHLHE40** | bHLH | Maturity_TF | SLOWER |
| **FOSL2** | BZIP, Fos | Maturity_TF | SHARP |
| **ZNF680** | C2H2 Kruepel KRAB | Maturity_TF | SHARP |
| **ZNF674** | C2H2 Kruepel KRAB | Maturity_TF | SHARP |
| **IRF4** | IRF | Maturity_TF | SHARP |
| **TGIF1** | homeo TALE/TGIF | Maturity_TF | SHARP |

| | | | |
|---|---|---|---|
| **ZFP64** | C2H2 Kruepel | Maturity_TF | SHARP |
| **ZNF561** | C2H2 Kruepel KRAB | Maturity_TF | SHARP |
| **NFKB1** | RHD | Maturity_TF | SHARP |
| **RXRA** | Nuclear hormone receptor | Maturity_TF | SHARP |
| **NUFIP1** | C2H2 | Maturity_TF | SHARP |
| **THRA** | Nuclear hormone receptor | Maturity_TF | SHARP |
| **POGK** | KRAB | Maturity_TF | SHARP |
| **PPARD** | Nuclear hormone receptor | Maturity_TF | SHARP |
| **ZNF189** | C2H2 Kruepel KRAB | Maturity_TF | SHARP |
| **HMGB2** | HMG box | Maturity_TF | SHARP |
| **ZNF526** | C2H2 Kruepel | Maturity_TF | SHARP |
| **CREB3L2** | BZIP, ATF | Maturity_TF | SLOWER |
| **ZNF557** | C2H2 Kruepel KRAB | Maturity_TF | SHARP |
| **ZHX1** | homeo ZHX | Maturity_TF | SHARP |
| **ZNF800** | C2H2 Kruepel | Maturity_TF | SHARP |
| **ZNF250** | C2H2 Kruepel KRAB | Maturity_TF | SHARP |
| **ZNF764** | C2H2 Kruepel KRAB | Maturity_TF | SHARP |
| **RELB** | RHD | Maturity_TF | SHARP |
| **ZFP82** | C2H2 Kruepel | Maturity_TF | SHARP |
| **NFIL3** | BZIP, NFIL3 | Maturity_TF | SHARP |
| **ZNF573** | C2H2 Kruepel KRAB | Maturity_TF | SHARP |
| **CREBL2** | BZIP, ATF | Maturity_TF | SLOWER |
| **ZNF568** | C2H2 Kruepel KRAB | Maturity_TF | SLOWER |
| **ZNF473** | C2H2 Kruepel KRAB | Maturity_TF | SLOWER |
| **ZNF331** | C2H2 Kruepel KRAB | Maturity_TF | SHARP |
| **ZNF354B** | C2H2 Kruepel | Maturity_TF | SHARP |
| **NR1H2** | Nuclear hormone receptor | Maturity_TF | SHARP |
| **ZNF41** | C2H2 Kruepel | Maturity_TF | SHARP |
| **ZNF404** | C2H2 Kruepel KRAB | Maturity_TF | SLOWER |
| **ZNF620** | C2H2 Kruepel KRAB | Maturity_TF | SHARP |
| **MYNN** | C2H2 Kruepel | Maturity_TF | SHARP |
| **ZNF138** | C2H2 Kruepel | Maturity_TF | SHARP |
| **ZKSCAN5** | C2H2 Kruepel KRAB | Maturity_TF | SHARP |
| **ZNF25** | C2H2 Kruepel | Maturity_TF | SHARP |
| **ZNF276** | C2H2 | Maturity_TF | SHARP |
| **NPAT** | NPAT | Maturity_TF | SHARP |
| **ATF1** | BZIP, ATF | Maturity_TF | SHARP |
| **ZNF793** | C2H2 Kruepel | Maturity_TF | SHARP |
| **ZHX3** | homeo ZHX | Maturity_TF | SHARP |
| **ZNF436** | C2H2 Kruepel KRAB | Maturity_TF | SHARP |
| **KLF6** | C2H2 Kruepel | Maturity_TF | SLOWER |
| **RORA** | Nuclear hormone receptor | Maturity_TF | SLOWER |
| **ZBTB16** | C2H2 Kruepel | Maturity_TF | SHARP |
| **MYC** | bHLH | Maturity_TF | SHARP |
| **STAT4** | STAT | Maturity_TF | SHARP |

| | | | |
|---|---|---|---|
| **NFATC2** | RHD | Maturity_TF | SHARP |
| **EOMES** | T-box | Maturity_TF | SHARP |
| **ZEB2** | Zinc finger E-box-binding homeobox | Maturity_TF | SHARP |
| **MAF** | BZIP, Maf | Maturity_TF | SHARP |
| **ZNF831** | C2H2 | Maturity_TF | SHARP |
| **HOPX** | homeo | Maturity_TF | SLOWER |
| **FOXO1** | Forkhead | Maturity_TF | SHARP |
| **TSC22D3** | TSC-22/Dip/Bun | Maturity_TF | SHARP |
| **RUNX3** | RUNX | Maturity_TF | SHARP |
| **IKZF3** | Ikaros C2H2 | Maturity_TF | SLOWER |
| **SKI** | SKI | Maturity_TF | SHARP |
| **SP140** | SAND | Maturity_TF | SHARP |
| **IRF1** | IRF | Maturity_TF | SHARP |
| **FOSB** | BZIP, Fos | Maturity_TF | SLOWER |
| **CEBPD** | BZIP, C/EBP | Maturity_TF | SHARP |
| **IRF7** | IRF | Maturity_TF | SHARP |
| **MYBL1** | MYB-HTH | Maturity_TF | SLOWER |
| **ZBTB38** | C2H2 ZBTB | Maturity_TF | SHARP |
| **ZNF766** | C2H2 Kruepel KRAB | Maturity_TF | SLOWER |
| **NR1D2** | Nuclear hormone receptor | Maturity_TF | SLOWER |
| **SMAD5** | MAD | Maturity_TF | SHARP |
| **ZBTB39** | C2H2 Kruepel | Maturity_TF | SHARP |
| **DACH1** | DACH | immaturity_related_TF | SLOWER |
| **FLI1** | ETS | immaturity_related_TF | SHARP |
| **ZNF415** | C2H2 | immaturity_related_TF | SHARP |
| **ZEB1** | Zinc finger E-box-binding homeobox | immaturity_related_TF | SHARP |
| **SMAD1** | MAD | immaturity_related_TF | SHARP |
| **ELF4** | ETS | immaturity_related_TF | SLOWER |
| **MEF2D** | MADS | immaturity_related_TF | SHARP |
| **ZNF596** | C2H2 Kruepel KRAB | immaturity_related_TF | SHARP |
| **ZNF92** | C2H2 Kruepel KRAB | immaturity_related_TF | SLOWER |
| **CSRNP1** | AXUD | immaturity_related_TF | SLOWER |
| **RORB** | Nuclear hormone receptor | immaturity_related_TF | SHARP |
| **KLF7** | C2H2 Kruepel | immaturity_related_TF | SLOWER |
| **ZNF251** | C2H2 Kruepel KRAB | immaturity_related_TF | SLOWER |
| **ZNF746** | C2H2 Kruepel KRAB | immaturity_related_TF | SLOWER |
| **NFYB** | NFY | immaturity_related_TF | SLOWER |
| **ADNP2** | C2H2 Kruepel | immaturity_related_TF | SLOWER |
| **MLXIP** | bHLH | immaturity_related_TF | SLOWER |
| **ZSCAN16** | C2H2 Kruepel | immaturity_related_TF | SLOWER |
| **ZNF175** | C2H2 Kruepel KRAB | immaturity_related_TF | SHARP |
| **ZNF275** | C2H2 Kruepel | immaturity_related_TF | SLOWER |
| **ZNF516** | C2H2 Kruepel | immaturity_related_TF | SHARP |
| **ZNF787** | C2H2 Kruepel | immaturity_related_TF | SHARP |
| **ZNF264** | C2H2 Kruepel KRAB | immaturity_related_TF | SHARP |

| | | | |
|---|---|---|---|
| **ZNF711** | C2H2 Kruepel | immaturity_related_TF | SLOWER |
| **NR4A3** | Nuclear hormone receptor | immaturity_related_TF | SLOWER |
| **ZNF627** | C2H2 Kruepel KRAB | immaturity_related_TF | SLOWER |
| **ZNF846** | C2H2 Kruepel | immaturity_related_TF | SLOWER |
| **ZNF470** | C2H2 Kruepel KRAB | immaturity_related_TF | SHARP |
| **PLAGL2** | C2H2 Kruepel | immaturity_related_TF | SLOWER |
| **MXD1** | bHLH | immaturity_related_TF | SHARP |
| **ZNF595** | C2H2 Kruepel KRAB | immaturity_related_TF | SHARP |
| **ZNF280C** | C2H2 | immaturity_related_TF | SHARP |
| **CREBZF** | BZIP, ATF | immaturity_related_TF | SLOWER |
| **ZNF670** | C2H2 Kruepel KRAB | immaturity_related_TF | SHARP |
| **ZNF506** | C2H2 Kruepel KRAB | immaturity_related_TF | SLOWER |
| **CTNND1** | Beta-catenin | immaturity_related_TF | SLOWER |
| **ZNF684** | C2H2 Kruepel KRAB | immaturity_related_TF | SLOWER |
| **ZNF570** | C2H2 Kruepel KRAB | immaturity_related_TF | SLOWER |
| **ZNF852** | C2H2 | immaturity_related_TF | SLOWER |
| **ZNF430** | C2H2 Kruepel KRAB | immaturity_related_TF | SHARP |
| **ZNF410** | C2H2 | immaturity_related_TF | SLOWER |
| **ZFP28** | C2H2 Kruepel | immaturity_related_TF | SHARP |
| **ZNF653** | C2H2 Kruepel | immaturity_related_TF | SHARP |
| **ZNF182** | C2H2 Kruepel KRAB | immaturity_related_TF | SLOWER |
| **ZNF790** | C2H2 Kruepel | immaturity_related_TF | SHARP |
| **ZNF571** | C2H2 Kruepel KRAB | immaturity_related_TF | SLOWER |
| **ZNF74** | C2H2 Kruepel | immaturity_related_TF | SHARP |
| **ZNF608** | C2H2 | immaturity_related_TF | SLOWER |
| **SOX4** | HMG SOX | immaturity_related_TF | SLOWER |
| **TCF7** | HMG LEF | immaturity_related_TF | SLOWER |
| **TOX2** | HMG box | immaturity_related_TF | SLOWER |
| **BACH2** | BZIP, CNC | immaturity_related_TF | SLOWER |
| **EGR2** | C2H2 EGR | immaturity_related_TF | SLOWER |
| **ZNF407** | C2H2 | immaturity_related_TF | SHARP |
| **HIVEP3** | C2H2 | immaturity_related_TF | SLOWER |
| **SATB1** | homeo cut | immaturity_related_TF | SLOWER |
| **RFX3** | RFX | immaturity_related_TF | SHARP |
| **LEF1** | HMG LEF | immaturity_related_TF | SLOWER |
| **ZNF236** | C2H2 Kruepel | immaturity_related_TF | SHARP |
| **ZBTB37** | C2H2 ZBTB | immaturity_related_TF | SLOWER |
| **TCF4** | bHLH | immaturity_related_TF | SHARP |
| **MYB** | MYB-HTH | immaturity_related_TF | SHARP |
| **ID3** | bHLH | immaturity_related_TF | SLOWER |
| **TSC22D1** | TSC-22/Dip/Bun | immaturity_related_TF | SLOWER |
| **ZNF280D** | C2H2 | immaturity_related_TF | SHARP |
| **MTA3** | GATA | immaturity_related_TF | SLOWER |
| **DBP** | BZIP, PAR | immaturity_related_TF | SHARP |
| **ZFX** | C2H2 Kruepel | immaturity_related_TF | SLOWER |

| | | | |
|---|---|---|---|
| **E2F5** | E2F/DP | immaturity_related_TF | SHARP |
| **TFDP2** | E2F/DP | immaturity_related_TF | SHARP |

**Supplementary table 11.** Demographic data of the Thymus samples.

| Batch | MAIT cells Stage | Sex /Age |
|---|---|---|
| **Thymus -1** | Stage 1 | Female/ 1 month 21 days |
| **Thymus -2** | Stage 1 | Female/ 4 months 17 days |
| **Thymus -3** | Stage 1, 2, 3 | Female/ 3 months 1 day |
| **Thymus -4** | Stage 1, 2, 3 | Female/ 4 years 8months |
| **Thymus -5** | Stage 2, 3 | Male/ 4 years |
| **Thymus -6** | Stage 2 | Male/ 7 days |
| **Thymus -7** | Stage 3 | Male/ 15 days |

**Supplementary table 1.** Antibody panel for defining human thymic MAIT cell stages 1-2.

| Immunogen | Conjugate | Manufacturer | Clone |
|---|---|---|---|
| | **Antibody stain- step 1 on ice** | | |
| **Biotinylated MR1** | Streptavidin PE-CF594 | BD Biosciences | |
| | **MACS Enrichment - step 2 on ice** | | |
| | **Antibody stain- step 3 on ice** | | |
| **CD4** | BV421 | BD Biosciences | RPA-T4 |
| **CD27** | BV510 | BD Biosciences | L128 |
| **CD1a** | BV786 | BD Biosciences | HI149 |
| **CD8** | BV605 | BD Biosciences | HIT8a |
| **CD161** | FITC | Miltenyi | 191B8 |
| **CD3** | PerCP/Cy5.5 | BD Biosciences | UCHT1 |

| | | | |
|---|---|---|---|
| **CD19** | APC-Cy7 | BioLegend | HIB19 |
| **CD14** | APC-Cy7 | BD Biosciences | MφP9 |
| **Va7.2** | BV711 | BioLegend | 3C10 |
| **Zombie NIR** | NIR | - | - |

**Supplementary table 2.** Antibody panel for defining human thymic MAIT cell stages 3.

| Immunogen | Conjugate | Manufacturer | Clone |
|---|---|---|---|
| **Antibody stain- step 1 on ice** | | | |
| **CD161** | PE -Vio770 | Miltenyi | 191B8 |
| **MACS Enrichment - step 2 on ice** | | | |
| **Antibody stain- step 3 on ice** | | | |
| **Biotinylated MR1** | Streptavidin PE-CF594 | BD Biosciences | - |
| **CD3** | PerCP/Cy5.5 | BD Biosciences | UCHT1 |
| **CD4** | BV421 | BD Biosciences | RPA-T4 |
| **CD8** | BV605 | BD Biosciences | HIT8a |
| **CD27** | BV510 | BD Biosciences | L128 |
| **CD19** | APC-Cy7 | BioLegend | HIB19 |
| **CD14** | APC-Cy7 | BD Biosciences | MφP9 |
| **Va7.2** | BV711 | BioLegend | 3C10 |
| **Zombie NIR** | NIR | - | - |

**Supplementary table 14.** General experimental buffers and reagents.

| Buffer components | Manufacturer | Cat number |
|---|---|---|
| **Molecular biology grade H2O (RNase, DNase free)** | VWR | VWRV02-0201-0500 |
| **NaCl** | Sigma-Aldrich | S5150-1L |

| | | |
|---|---|---|
| **Digitonin** | Millipore Sigma | 300410 |
| **Trypan Blue** | Thermo Fisher Scientific | T10282 |
| **Spermidine trihydrochloride*** | Sigma-Aldrich | S0266 |
| **Triton X-100** | Sigma-Aldrich | X100 |
| **EDTA (prepare 0.5 M stock at pH 8.0)** | Sigma-Aldrich | E5134 |
| **complete$^{TM}$, Mini, EDTA-free Protease Inhibitor Cocktail** | Roche | 11836170001 |
| **HEPES** | Sigma-Aldrich | H3375 |
| **DMSO** | Sigma | D8418-100ml |
| **Glycerol** | Millipore | G5516 |
| **Sodium dodecyl sulfate (SDS)** | Sigma-Aldrich | L4509 |
| **MnCl2** | Sigma-Aldrich | 203734 |
| **KCl** | Sigma-Aldrich | P3911 |
| **1 M TAPS, pH8.5** | Sigma | T5130 |
| **CaCl2** | Sigma-Aldrich | C1016 |

**Supplementary table 15.** Reagents for the CUT&Tag assay.

| Cut &Tag Reagent | Manufacturer | Cat number |
|---|---|---|
| **Concanavalin A (ConA) Conjugated Paramagnetic Beads** | EpiCypher | 21-1401 |
| **CUTANA$^{TM}$ pAG-Tn5** | EpiCypher | 15-1017,15-1117 |
| **Antibody to histone PTM** | Diagenode | C15410196 |
| **Anti-Rabbit Secondary Antibody** | EpiCypher | 13-0047 |
| **Agencourt AMPure XP magnetic beads** | Beckman Coulter | A63880 |
| **Qubit$^{TM}$ 1x dsDNA HS Assay Kit** | Thermo Fisher Scientific | Q33230 |
| **CUTANA High-Fidelity 2X PCR Master Mix$^{TM}$** | EpiCypher | 15-1018 |
| **Universal i5 Primer** | IDT | |
| **Uniquely Barcoded i7 Primers** | IDT | |
| **Rabbit IgG Negative Control Antibody** | EpiCypher | 13-0042 |

**Supplementary table 16.** CUT&Tag buffer compositions (adapted from EpiCypher protocol v2.0).

| **Buffers (preparation)** | |
|---|---|
| **Nuclear Extraction (NE) Buffer** | 20 mM HEPES-KOH (pH 7.9), 10 mM KCl, 0.1% Triton X-100, 20% glycerol, 0.5 mM spermidine, 1× cOmplete™ Mini EDTA-free protease inhibitor |
| **Bead Activation Buffer** | 20 mM HEPES (pH 7.9), 10 mM KCl, 1 mM $CaCl_2$, 1 mM $MnCl_2$ |
| **Wash150 Buffer** | 20 mM HEPES (pH 7.5), 150 mM NaCl, 0.5 mM spermidine, 1× cOmplete™ Mini EDTA-free protease inhibitor |
| **Digitonin150 Buffer** | Wash300 buffer, 0.01% digitonin |
| **Antibody150 Buffer** | Digitonin150 buffer, 2 mM EDTA |
| **Wash300 Buffer** | 20 mM HEPES (pH 7.5), 300 mM NaCl, 0.5 mM spermidine, 1× cOmplete™ Mini EDTA-free protease inhibitor |
| **Digitonin300 Buffer** | Wash300 buffer, 0.01% digitonin |
| **Tagmentation Buffer** | Digitonin300 buffer, 10 mM $MgCl_2$ |
| **TAPS Buffer** | 10 mM TAPS (pH 8.5), 0.2 mM EDTA |
| **SDS Release Buffer** | 10 mM TAPS (pH 8.5), 0.1% SDS |
| **SDS Quench Buffer** | 0.67% Triton X-100 in molecular biology–grade $H_2O$ |

Buffer compositions were adapted from the EpiCypher CUT&Tag protocol (v2.0, revised 7 August 2024).
